# Proteomics of autophagy deficient macrophages reveals enhanced antimicrobial immunity via the oxidative stress response

**DOI:** 10.1101/2020.09.10.291344

**Authors:** Timurs Maculins, Erik Verschueren, Trent Hinkle, Patrick Chang, Cecile Chalouni, Junghyun Lim, Anand Kumar Katakam, Ryan C. Kunz, Brian K. Erickson, Ting Huang, Meena Choi, Tsung-Heng Tsai, Olga Vitek, Mike Reichelt, John Rohde, Ivan Dikic, Donald S. Kirkpatrick, Aditya Murthy

## Abstract

Defective autophagy is associated with chronic inflammation. Loss-of-function of the core autophagy gene Atg16l1 increases risk for Crohn’s disease by enhancing innate immunity in macrophages. However, autophagy also mediates clearance of intracellular pathogens. These divergent observations prompted a re-evaluation of ATG16L1 in antimicrobial immunity. In this study, we found that loss of Atg16l1 in macrophages enhanced the killing of virulent *Shigella flexneri* (*S.flexneri*), an enteric bacterium that resides within the cytosol by escaping all membrane-bound compartments. Quantitative multiplexed proteomics revealed that ATG16L1 deficiency significantly upregulated proteins involved in the glutathione-mediated antioxidant response to compensate for elevated oxidative stress, which also promoted *S.flexneri* killing. Consistently, myeloid cell-specific deletion of Atg16l1 accelerated bacterial clearance *in vivo*. Finally, pharmacological modulation of oxidative stress by suppression of cysteine import conferred enhanced microbicidal properties to wild type macrophages. These findings demonstrate that control of oxidative stress by ATG16L1 regulates antimicrobial immunity against intracellular pathogens.

**Impact statement:** Maculins *et al* utilize multiplexed mass spectrometry to show that loss of the autophagy gene *Atg16l1* in macrophages enhances antimicrobial immunity against intracellular pathogens via the oxidative stress response.

## Introduction

Effective immunity against enteric pathogens requires complex signaling to coordinate the inflammatory response, pathogen clearance, tissue remodeling and repair (Maloy and Powrie, 2011). Autophagy, a cellular catabolic pathway that eliminates cytosolic cargo via lysosomal degradation, has emerged as an important regulator of mucosal immunity and inflammatory bowel disease (IBD) etiology. Genome-wide association studies linked a missense variant (T300A) in the core autophagy gene *Atg16l1* with increased risk for inflammatory bowel diseases (Hampe et al., 2007; Rioux et al., 2007). Later studies demonstrated that this variant contributes to enhanced caspase-mediated degradation of the ATG16L1 protein (Lassen et al., 2014; Murthy et al., 2014). Genetic loss-of-function of core autophagy genes including *Atg16l1* increases secretion of pro-inflammatory cytokines by macrophages in response to toll-like receptor (TLR) activation (Lim et al., 2019; Saitoh et al., 2008). This contributes to increased mucosal inflammation, driving resistance to extracellular pathogens such as *Citrobacter rodentium* (Marchiando et al., 2013; Martin et al., 2018) and pathogenic *Escherichia coli* (Wang et al., 2019). Defective autophagy in the myeloid compartment also confers enhanced antimicrobial immunity against certain intracellular pathogens, such as *Salmonella typhimurium* (*S.typhimurium*) and *Listeria monocytogenes* via induction of type I and II interferon responses (Samie et al., 2018; Wang et al., 2020). Thus, autophagy acts as an immuno-suppressive pathway in antimicrobial immunity *in vivo*.

Targeted elimination of intracellular pathogens by xenophagy, a form of selective autophagy, is well-described in cellular model systems (Bauckman et al., 2015). In contrast to non-selective autophagy triggered by nutrient stress, xenophagy functions to eliminate intracellular bacteria by sequestering them in autophagosomes and shuttling them to the degradative lysosomal compartment. Pathogenic bacteria have evolved mechanisms to either evade capture by the autophagy machinery, as by *S.typhimurium* and *Shigella flexneri* (*S.flexneri*) (Birmingham et al., 2006; Campbell-Valois et al., 2015; Dong et al., 2012; Martin et al., 2018; Xu et al., 2019b) or attenuate autophagic flux as by *Legionella pneumophila* (Choy et al., 2012). *S.typhimurium* primarily resides in a protective compartment known as the *Salmonella* containing vacuole (SCV). There it prevents formation of the ATG5-ATG12-ATG16L1 complex at the bacterial vacuolar membrane via secretion of the effector SopF, which blocks ATG16L1 association with vacuolar ATPases (Xu et al., 2019b). Despite its ability to interfere with autophagy, infected host cells still recognize 10-20% of cytosolic *S.typhimurium* and subject this sub-population to lysosomal degradation via mechanisms involving direct recognition of either the bacterial surface (Huang and Brumell, 2014; Stolz et al., 2014) or damaged phagocytic membranes (Fujita et al., 2013; Thurston et al., 2012).

Compared to *S.typhimurium, S.flexneri* is not characterized by a vacuolar life cycle, but instead resides in the host cytoplasm. *S.flexneri* effector proteins IcsB and VirA are capable of completely inhibiting autophagic recognition to permit replication in the host cytosol (Liu et al., 2018; Ogawa et al., 2005). In response, the host cell attempts to further counteract *S.flexneri* infection via diverse mechanisms, such as coating bacterial cell surfaces with guanylate-binding proteins (GBPs) (Li et al., 2017; Wandel et al., 2017) or sequestering bacteria in septin cage-like structures to restrict their motility (Mostowy et al., 2010). To reveal these mechanisms, cell-based studies have largely utilized attenuated variants (e.g. IcsB or IcsB/VirA double mutants of *S.flexneri*) or strains that inefficiently colonize the host cytosol (e.g. *S.typhimurium* which express SopF). Thus, observations from *in vivo* genetic models must be reconciled with observations made in cell-based systems to fully describe the roles of autophagy in antimicrobial immunity. Importantly, there is a lack of understanding of how autophagy contributes to immunity against non-attenuated (wild type) cytosolic pathogens. This insight is especially lacking in relevant cell types, such as macrophages that constitute a physiologically relevant niche for the expansion of *S.flexneri* (Ashida et al., 2015).

In this study we investigated the role of macrophage ATG16L1 in response to infection by wild type *S.flexneri* (strain M90T). Surprisingly, we observed that loss of *Atg16l1* in BMDMs enhanced *S.flexneri* elimination in culture, as well as by mice lacking ATG16L1 in the myeloid compartment *in vivo* (*Atg16l1-cKO*). We utilized multiplexed quantitative proteomics to characterize total protein, phosphorylation and ubiquitination changes in wild type (WT) and ATG16L1-deficient (cKO) bone marrow-derived macrophages (BMDMs) either uninfected or infected with *S.flexneri*. Quantifying global protein levels along with site-specific post-translational modifications (PTMs) provided a comprehensive catalogue of basal differences between WT and cKO BMDMs and the dynamic response of each to infection. As expected, profound differences were observed for components in the autophagy pathway, as well as proteins involved in cell death, innate immune sensing and NF-κB signaling. Interestingly, a cluster of proteins emerging from the proteomics data implicated the basal oxidative stress response as a key difference between control and ATG16L1-deficient BMDMs. In particular, significant accumulation of the SLC7A11 subunit of a sodium-independent cystine*-*glutamate antiporter (XCT), critical for the generation of glutathione (GSH) used in detoxification of ROS and lipid peroxides, was noteworthy in cKO cells. This coincided with basal elevation of cytosolic ROS in cKO BMDMs, thus providing an explanation for the sustained viability and antimicrobial capacity of ATG16L1-deficient macrophages. Furthermore, increased cytosolic ROS caused by pharmacological XCT inhibition enhanced *S.flexneri* clearance by WT BMDMs, recapitulating cKO phenotypes. Taken together, this study offers a comprehensive, multidimensional catalogue of proteome-wide changes in macrophages following infection by an enteric cytosolic pathogen, including key nodes of cell-autonomous immunity regulated by autophagy. Our findings demonstrate that ATG16L1 tunes antimicrobial immunity against cytosolic pathogens via the oxidative stress response, and that pharmacological modulation of this pathway represents a novel strategy towards enhanced elimination of cytosolic pathogens.

## Results

### Enhanced clearance of intracellular *S.flexneri* by loss of *Atg16l1*

Recent studies have identified that defective autophagy in macrophages enhances multiple inflammatory signaling responses to promote antimicrobial immunity (Lim et al., 2019; Martin et al., 2018; Samie et al., 2018; Wang et al., 2020). Given these observations, we wanted to explore whether loss of *Atg16l1* affects killing of the wild type, invasive, intracellular pathogen *Shigella flexneri* strain M90T (*S.flexneri*). To test this, bone marrow-derived macrophages (BMDMs) from either control (*Atg16l1-WT*) or mice lacking ATG16L1 in the myeloid compartment (*Atg16l1-cKO*) were subjected to the gentamycin protection assay that enables quantification of intracellular bacteria by enumerating colony forming units (CFUs). We first determined the kinetics of *S.flexneri* killing by following BMDM infection over six hours (MOI 5). Compared to wild type (WT) controls, ATG16L1-deficient BMDMs (cKO) demonstrated accelerated bacterial clearance starting at two hours post-infection (Figure 1A and 1B). Previous studies demonstrated enhanced sensitivity of autophagy-deficient cells to programmed cell death following engagement of cytokine receptors and microbial ligands (Lim et al., 2019; Matsuzawa-Ishimoto et al., 2017; Orvedahl et al., 2019). Thus, BMDM viability was measured in parallel by quantifying the propidium iodide (PI)-positive population via live-cell imaging. WT and cKO BMDMs displayed similar cell death kinetics over the time course of infection, indicating that accelerated *S.flexneri* killing was not driven by enhanced cell death, but potentially by other cytosolic factors in cKO BMDMs (Figure 1C).

**Figure 1.**
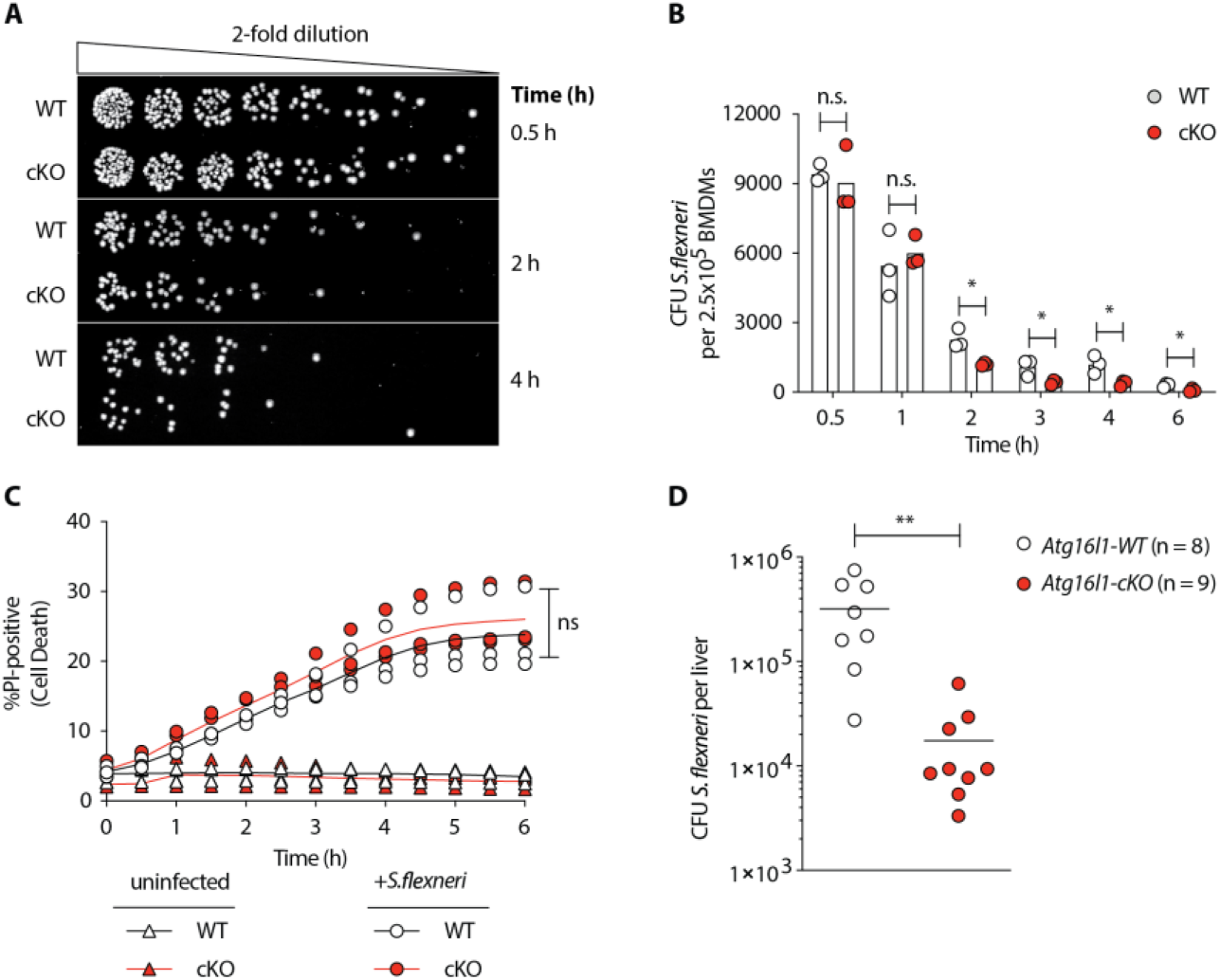
Enhanced clearance of intracellular *S.flexneri by* loss *of Atg16l1*. (A) Representative serial dilutions from gentamycin protection assays following *S.flexneri* M90T infection of WT or cKO BMDMs at the indicated timepoints. (B) Comparison of colony forming units (CFUs) per well from three independent infection experiments using BMDM preparations from three different *Atg16H-WT* or *Atg16l1-cKO* mice, ns, non-significant; 2h * P = 0.01,3h * P = 0.03, 4h * P = 0.02, 6h * P = 0.03, multiple t-test comparison. (C) Percentage of propidium iodide (Pl)-positive cells during time-course infection of WT or cKO BMDMs with *S.flexneri* M90T. Graph represents individual values from three independent experiments using three different BMDM preparations, ns, non-significant. (D) Liver bacterial load 24 hours following intravenous injection of *Atg16H-WT* or *Atg16H-cKO* mice with *S.flexneri* M90T. Graph shows data from a representative experiment out of four different experiments as Iog10 CFU count per liver in indicated number of mice, **P = 0.0031. Outliers removed using ROUT (Q = 1%) method.

To corroborate this finding *in vivo*, control and *Atg16l1-cKO* mice were infected with the *S.flexneri* via tail vein injection and CFUs enumerated from hepatic lysates. Myeloid-specific loss of *Atg16l1* resulted in a markedly decreased bacterial burden 24 hours post-infection (Figure 1D). Taken together, these observations establish that ATG16L1 restrains macrophage immunity against cytosolic bacteria such as *S.flexneri*.

### Multiplexed proteomic profiling of autophagy competent and deficient BMDMs following infection

To reveal factors that may drive enhanced *S.flexneri* killing in ATG16L1-deficient BMDMs, we characterized changes in global proteome and post-translational modifications (PTMs) in proteins between WT and cKO BMDMs. To that end, we applied tandem mass spectrometry coupled with tandem mass tagging (TMT) 11-plex isobaric multiplexing. Cell lysates were prepared from WT and cKO BMDMs that were either uninfected (U) or infected at early (E; 45-60min) or late (L; 3-3.5h) time-points with *S.flexneri* (MOI 5). Cumulatively, two 11-plex experiments were performed with uninfected samples represented in biological triplicates and infected samples represented in biological quadruplicates (see Methods for details) (Figure 2A). Data were acquired using the recently established SPS-MS3 approach wherein dedicated MS3 scan events are collected from fragment ion populations representing a mixture of the 11 samples and used to report the relative abundance of each peptide feature per channel (McAlister et al., 2014; Ting et al., 2011).

**Figure 2.**
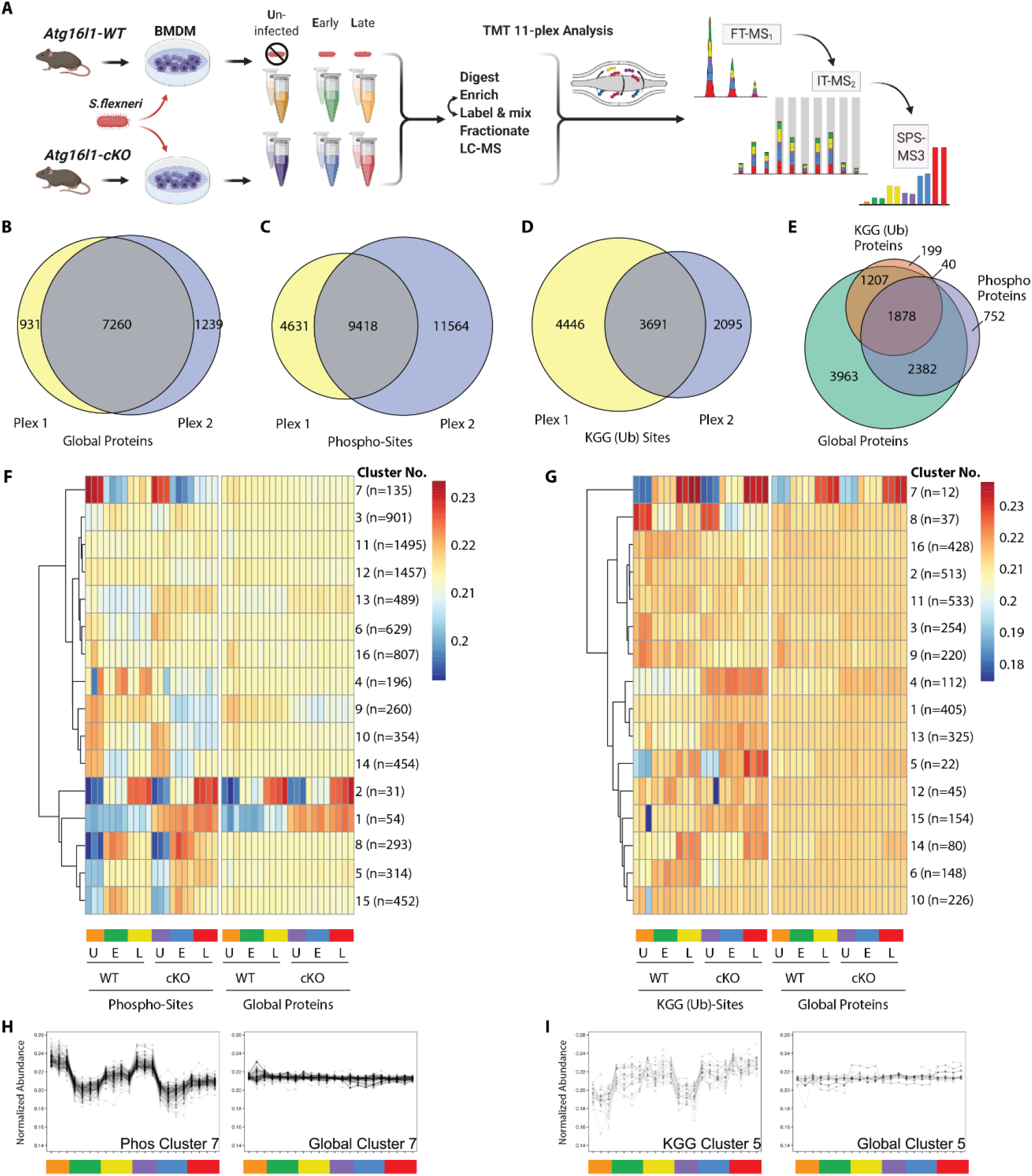
Multiplexed proteomic profiling of autophagy competent and deficient BMDMs following infection. (A) Schematic representation of multiplexed proteomic profiling of macrophages during *S.flexneri* infection. (B-D) Venn diagrams show overlapping quantitative data collected in Plexl and/or Plex2 for (B) Global Proteins, (C) Phosphorylation sites and (D) KGG (Ub) sites. (E) Venn diagram displays an overlap of quantitative data for Phospho- and KGG (Ub) sites with respect to the Global Proteins quantified. (F and G) Heatmaps displaying K-means clustered quantitative data for (F) Phospho-sites and (G) KGG (Ub) sites relative to their corresponding Global Proteins. Note that Global Protein clustering differs between panels F and G based on the proteins from which PTMs were quantified. (H and I) Line plots showing representative clusters from the Heatmaps above. Phospho Cluster 7 (panel H) and KGG (Ub) Cluster 5 (panel I) each show PTM profiles that diverge from their corresponding Global Protein measurements. Proteins and PTMs making up each cluster are presented in Table S1.

For global proteome profiling, quantitative data was obtained from >103,700 unique peptides mapping to 9430 proteins. From the PTM enriched samples, quantitative data were obtained for >25,600 unique phosphorylation sites (5052 proteins) and >12,400 unique KGG (Ub) sites (3324 proteins). When considering only features bearing data in both 11-plexes, the final dataset contained 22 channels of quantitative data for 7260 proteins (i.e. global proteome), 9418 phosphorylation sites and 3691 KGG modification sites (Figure 2B-D). As expected, ~90% of the post-translationally modified peptide spectral matches derived from proteins that were also identified and quantified in the global proteome dataset (Figure 2E). Both within and between plexes, peptide and protein level quantitative data were highly reproducible with Pearson correlations ranging from 0.96-0.99 (Figure S1A). Phosphorylation and KGG profiling data were subjected to K-means clustering, each paired with the corresponding global proteome data. Heatmap representations revealed clusters of PTM changes that occur in genotype and/or infection dependent manners (Figure 2F and 2G). A subset of these clusters comprised PTMs whose quantitative profiles mirrored that of the underlying protein level due to altered protein expression or stability (e.g. Phospho Clusters 1-2 in Figure 2F and S1B; KGG Cluster 7 in Figure 2G and S1C). In contrast, other clusters displayed PTM profiles that diverged from their underlying proteins (e.g. Phospho Cluster 7 in Figure 2F and 2H; KGG Cluster 5 in Figure 2G and 2I). The composition of PTMs and proteins comprising each cluster are available in Table S1.

Interrogation of the uninfected datasets revealed differences between the genotypes on the global protein level. Consistent with previous observations (Samie et al., 2018), cKO BMDMs showed upregulation in autophagy receptors, such as SQSTM/p62 and ZBP1 (Figure S2A). In the phosphorylation and KGG datasets, interesting observations amongst others concerned elevated phosphorylation of ubiquitin (RL40) at serine (S) 57 and ubiquitination of FIS1 at lysine (K) 20, which are involved in endocytic trafficking (Lee et al., 2017; Peng et al., 2003) and mitochondrial and peroxisomal homeostasis (Bingol et al., 2014; Koch et al., 2005; Zhang et al., 2012), respectively (Figure S2B and S2C).

Interrogation of the infected datasets revealed the dynamic nature of the macrophage response to infection. For example, global proteome analysis revealed broad changes in pro-inflammatory cytokines and chemokines at early (GROA), late (CXL10, IL1A, IL1B) or both (CCL2, TNFA) time-points, as well as marked changes in several key cell surface receptors (Figure S2D, S3A and S3B). Time-dependent changes were also observed for components of innate immune signaling that intersect with the ubiquitin pathway (PELI1), kinase-phosphatase signaling (DUS1/Dusp1) and GTP/GDP signaling (GBP5) (Figure S3C). For phosphorylation, notable examples included tyrosine (Y) 431 of the PI3-kinase regulatory subunit (P85A) and S379 of the interferon regulatory factor (IRF3) (Figure S2E). In the case of ubiquitination, marked effects are seen for a selective autophagy receptor Tax1BP1 (TAXB1_K618) and an E3 ubiquitin ligase Pellino (PELI1_K202) (Figure S2F), both of which have defined roles at the intersection of cell death and innate immune signaling (Choi et al., 2018; Gao et al., 2011; Parvatiyar et al., 2010).

It is beyond of the scope of this study to describe these in-depth proteomic observations. Therefore, we developed interactive Spotfire Dashboards as a resource to facilitate discoveries in cellular pathways of interest by other investigators. These can be accessed at the following URL: https://info.perkinelmer.com/analytics-resource-center.

### Characterizing PTMs of autophagy proteins and inflammatory signaling nodes revealed by loss of *Atg16l1* and infection

To effectively integrate data for each protein within a single consolidated view, heatmaps were assembled to show the proteome level change immediately adjacent to any PTMs that were identified in the phospho- and KGG-enriched samples. In the example for selective autophagy receptor Tax1bp1 (TAXB1), heatmaps depict relative abundance of features present in one or both experiments (Plex1 and/or Plex2) (Figure 3A). Comparisons of interest include cKO versus WT for uninfected, early and late infection time-point samples. For TAXB1, these show that the global protein level is elevated upon *Atg16l1* deletion, as are a number of individual phosphorylation and ubiquitination sites including those quantified in one (e.g. T494, K624) or both plexes (e.g. S632, S693, K627). Additional comparisons call out time-dependent differences between infected and uninfected conditions for each genotype - namely early versus uninfected (E/U) and late versus uninfected (L/U). For TAXB1, certain PTMs such as phosphorylation at S632 and ubiquitination at K624 and K627 track with the protein, while other PTMs such as phosphorylation at threonine (T) 494 and S693 display time-dependent changes that diverge from the underlying protein level (Figure 3A). Shown individually, histograms depict the same information for relative abundance of TAXB1 and its specific PTMs (Figure 3B). Therefore, the heatmaps provide a succinct visual representation of all detected changes in protein and PTM abundance.

**Figure 3.**
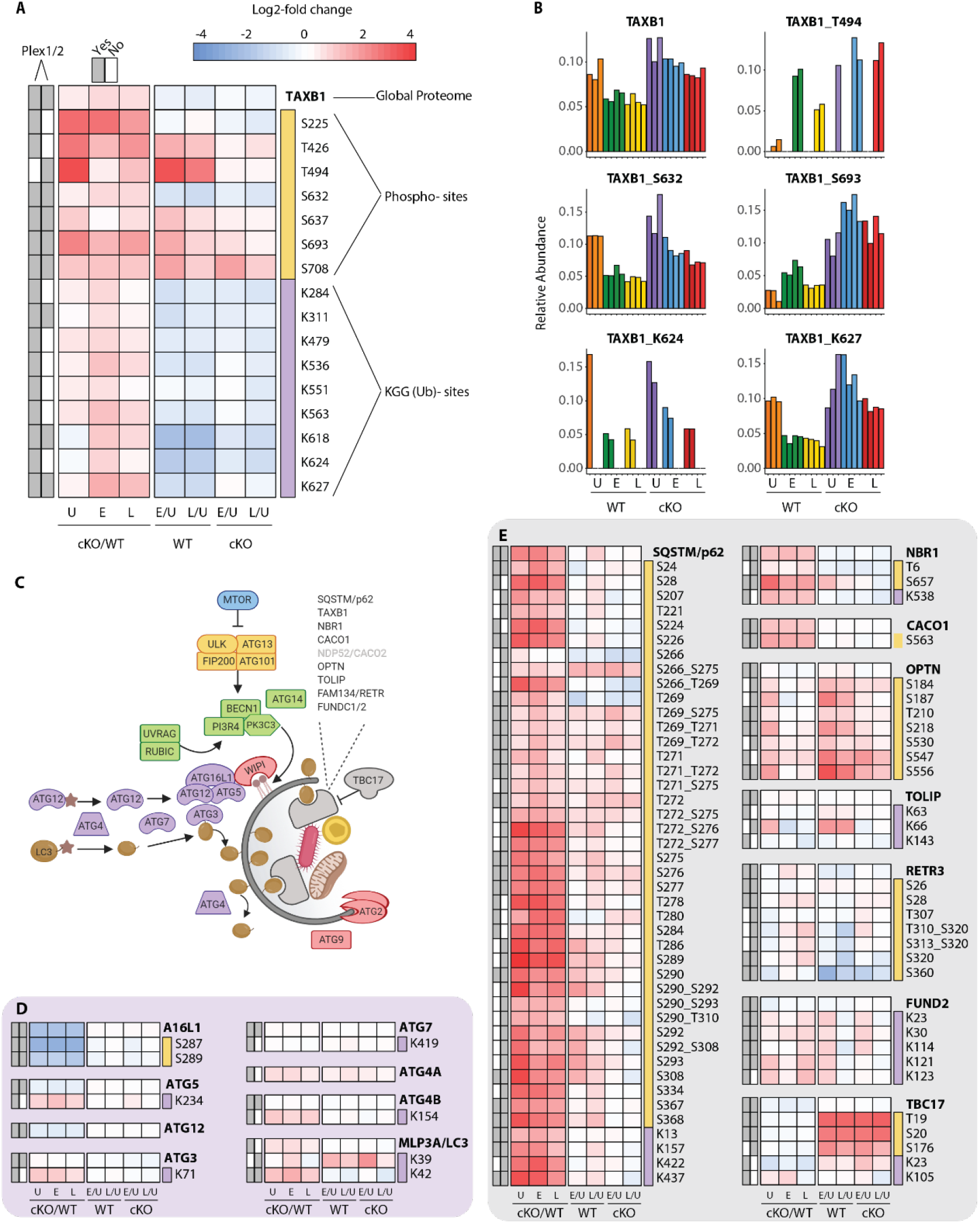
Characterization of proteomic changes in the autophagy pathway. (A) Heatmap representation of log2 fold changes for global proteome (unmarked), phospho-(yellow section) or KGG (Ub)- sites (purple section) measurements made forTAXBI. Data are shown for features quantified from uninfected (U) WT and cKO BMDMs or cells infected at early (E) or late (L) timepoints with *S.flexneri*. Log2 transformed ratios are shown for contrasting genotypes (cKO/WT) at each infection timepoint (U, E, L) on the left and between infection timepoints (E/U and L/U) within each genotype on the right. Grey boxes denote quantification of the feature in Plexl and/or Plex2. Modification sites on TAXB1 denote the modified amino acid (S/T/Y/K) and residue number. (B) Bar graphs showing the relative abundance of TAXB1 global protein and representative phospho- and KGG (Ub)- sites in each of the six conditions. Note that TAXBI_K624 (Plexl) and TAXBI_T494 (Plex2) represent data collected only in a single Plex, with the relative abundance of TMT reporter ions summing up to 1.0. (C) Schematic representation of macro-autophagy & selective autophagy machinery. (D and E) Heatmap representations of E1 /E2/E3-like pathway components responsible for conjugating LC3 (MLP3A) to regulate autophagosome membrane elongation (D) and selective autophagy receptors (E). The background shading for each panel corresponds to the functional color coding of proteins in the pathway schematic shown in (C).

One pathway where we expected to see marked proteome and PTM level changes upon infection was in autophagy (Figure 3C and S4). We confirmed genotype-dependent effects on each component of the ATG5-ATG12-ATG16L1 E3 ligase-like complex that conjugates LC3 (MLP3A) to phosphatidylethanolamine (Figure 3D). Only modest changes were seen in the core autophagy machinery following infection, with the most notable effects being differential phosphorylation of FIP200 (RBCC1), ATG2B, and VPS15/p150 (PI3R4) (Figure S4C-E). More substantial effects were seen for phosphorylation events on autophagy receptors such SQSTM/p62, Optineurin (OPTN) (Figure 3E) and TAXB1 (Figure 3A). In the case of p62, singly and multiply phosphorylated forms of T269, T271, T272, S275/6, S277 were elevated in ATG16L1-deficient macrophages, most notably at the early timepoint post-infection. S28 phosphorylation of p62 was previously described to regulate activation of the antioxidant response (Xu et al., 2019a). Interestingly, we detected a substantial increase in basal S28 phosphorylation in cKO BMDMs, indicating that ATG16L1 deficiency may impact oxidative stress (Figure S4F).

Our PTM datasets showed dynamic regulation of a range of inflammatory signaling components by infection as well as autophagy (Figure S5). For example, we detected ubiquitination of K278 of NEMO (Figure S5F), consistent with increased LUBAC activity (Tokunaga et al., 2009). Interestingly, the global proteome data reported a peptide with the sequence GGMQIFVK that is derived from linear polyubiquitin chains formed by the LUBAC complex. This linear ubiquitin peptide was elevated upon infection in both WT and cKO BMDMs (Figure S3D), further supporting increased E3 ubiquitin ligase activity of LUBAC. As noted above, TAXB1 phosphorylation was induced upon infection at a number of sites (Figure 3A). These changes in TAXB1 correlated with numerous elevated PTMs of the A20 (TNAP3) deubiquitinase, a protein whose anti-inflammatory activity modulates NF-κB signaling (Figure S5C). Interestingly, phosphorylation at S693 of TAXB1 is important for the assembly of TNAP3-containing complex and negative regulation of NF-κB signaling (Shembade et al., 2011) (Figure 3A).

We also identified notable changes across numerous components implicated in pathogen sensing such as TLRs, RLRs, NLRs and STING/cGAS (Supplementary Figure S6A and S6B). Our datasets confirm numerous previously demonstrated PTMs that occur in response to infection, such as elevated phosphorylation of RIPK1 at S321 (Figure S5E), XIAP at S429 or IRF3 on multiple sites (Figure S6D and S6E). Similar effects were observed for ABIN1 (TNIP1), which showed minimal changes in global protein levels, but elevated ubiquitination at multiple lysines including K360, K402, K480 at both timepoints and higher levels in cKO than WT (Figure S5F). Caspase-8 ubiquitination was elevated at K169 in both WT and cKO early post-infection, but was sustained through the late timepoint only in ATG16L1-deficient BMDMs (Figure S5G). Within the ubiquitin pathway, E3 ubiquitin ligases including HOIP (RNF31), TRAF2, and Pellino (PELI1) showed marked infection dependent changes at the level of phosphorylation (e.g. RNF31_S445) and ubiquitination (e.g. PELI_K202 early, TRAF2_K313 late) (Figure S5C).

Cross-referencing all highlighted PTMs with PhosphoSitePlus^®^ revealed that ~60% of PTMs were previously identified in distinct large-scale proteomic screens without assigning a specific biological role, but only 15% of PTMs have been studied in connection to a biological function (Tables S2 and S3). This analysis also revealed that nearly 25% of PTMs in autophagy, innate sensing, inflammatory and cell death signaling identified in our study appear to be novel (summarized in Table 1).

**Table 1.**
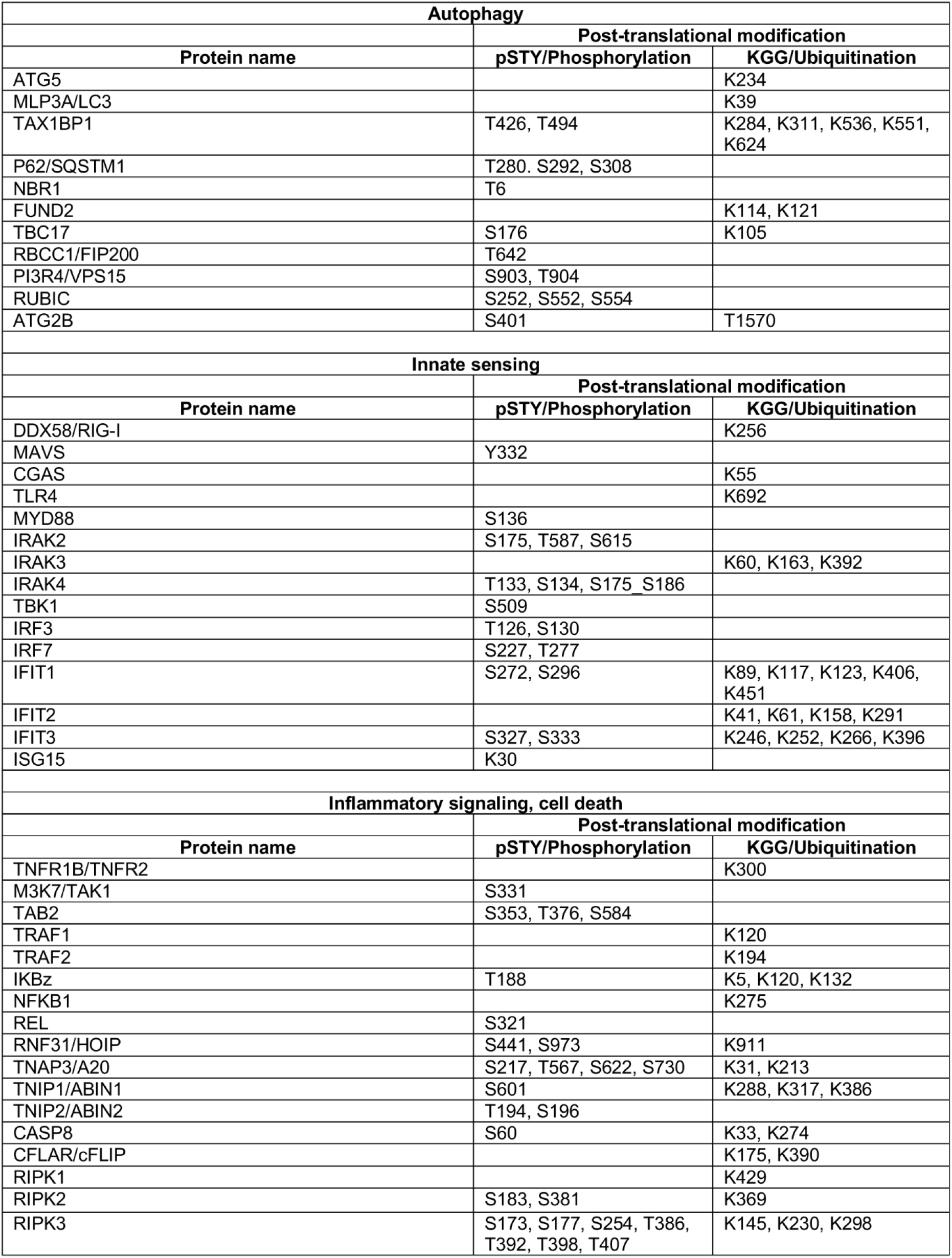
Novel post-translational modifications in specific autophagy, innate sensing, inflammatory signaling and cell death pathways revealed by TMT-MS of BMDMs following *S.flexneri* infection.

### Elevated oxidative stress in ATG16L1-deficient macrophages contributes to accelerated bacterial killing

To reveal cellular processes overrepresented in cKO BMDMs in an unbiased manner, we leveraged the deep coverage of the global proteome by TMT-MS and performed gene set enrichment analysis (GSEA). Unexpectedly, this analysis revealed a strong enrichment of the components of reactive oxygen species (ROS) pathway (Figure 4A and Table S4 for protein set terms). Further assessment of the components of this gene ontology term revealed critical regulators of redox homeostasis that were increased in uninfected cKO BMDMs at steady state relative to WT (Figure 4B). This group of proteins included several factors involved in glutathione (GSH) synthesis, such as the glutamate-cysteine ligase regulatory subunit (GSH0/Glcm) and GSH synthetase (GSHB/Gss), and GSH regeneration, such as microsomal glutathione S-transferase (MGST1) and NAD(P)H dehydrogenase 1 (NQO1). Additionally, several ROS converting enzymes including catalase (CATA) and peroxiredoxin 1 (PRDX1) were also elevated in cKO BMDMs at steady state. Furthermore, a subset of these redox regulators changed abundance upon *S.flexneri* infection. For example, prostaglandin dehydrogenase 1 (PGDH) displayed a time dependent decrease upon infection that was accentuated in cKO versus WT, consistent with its known susceptibility to ROS (Wang et al., 2018). Conversely, levels of the cysteine-glutamate antiporter SLC7A11 (XCT) (Conrad and Sato, 2012; Sato et al., 1999) exhibited a significant increase in cKO BMDMs following infection (Figure 4C). Thus, ATG16L1 deficiency and *S. flexneri* infection might each independently elevate ROS levels, with ATG16L1 deficiency further driving a compensatory increase in the lipid ROS regulatory pathway during infection to maintain macrophage viability.

**Figure 4.**
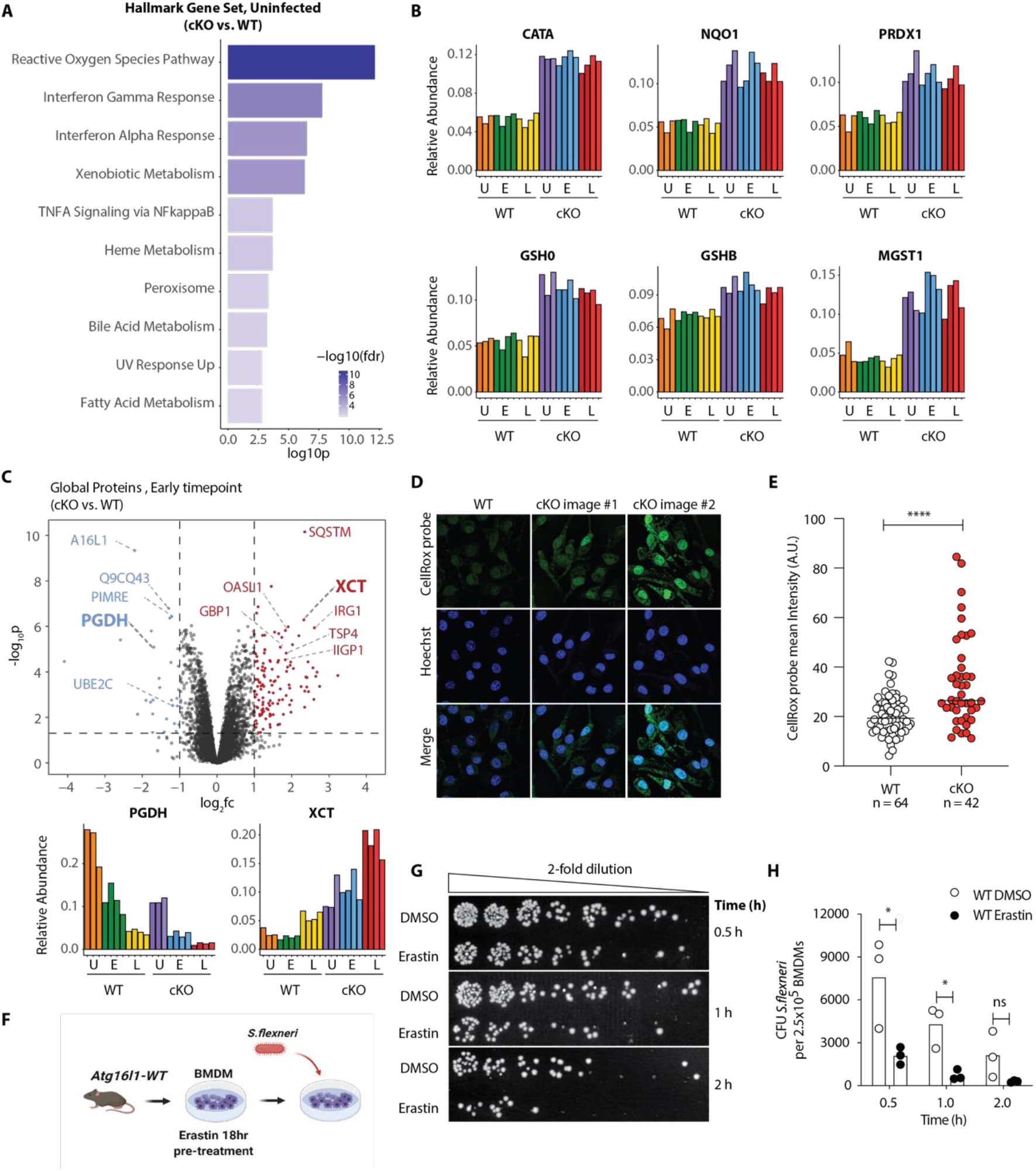
Elevated oxidative stress in ATG16L1 -deficient macrophages contributes to accelerated bacterial killing. (A) Gene set enrichment analysis (GSEA) of global proteome data showing cellular processes overrepresented in uninfected cKO over WT BMDMs. (B) Bar graphs showing the relative abundances for selected proteins involved in redox regulation and detoxifying reactive oxygen species. (C) Volcano plot of global protein changes at early infection timepoint between the genotypes. Proteins enriched in cKO and WT BMDMs are highlighted in red and blue, respec-tively. Bar graphs showing the cumulative effects of genotype and infection on PGDH and XCT protein levels. (D) Representative images from experiments shown in (E) demonstrating CellRox probe intensity, Hoechst nuclear staining and merged images. (E) Quantification of CellRox green mean intensity in WT and cKO BMDMs. Graph shows single cell data from a representative experiment (n = 3). Unpaired t test **” P < 0.0001. (F) Schematic representa-tion of infection experiment using pre-treatment of BMDMs with Erastin. (G) Representative serial dilutions from gentamycin protection assays following *S.flexneri* M9OT infection of WT BMDMs in the presence of DMSO or Erastin (4 μg/ml) at the indicated timepoints. Erastin-treated WT BMDMs were pre-treated with Erastin for 18 hours prior infection (T = 0). (H) Comparison of CFUs from three independent infection experiments using BMDM preparations from three different *Atg16l1-WT* mice, ns, non-significant; 0.5 h * P = 0.04,1 h * P = 0.01.

To determine if ATG16L1-deficient BMDMs are exposed to higher oxidative stress, we first used a fluorogenic probe (CellRox green) that enables measurement of oxidative stress by confocal fluorescence microscopy. Interestingly, despite upregulation of numerous redox regulatory factors, CellRox probe intensity was significantly higher in cKO BMDMs (Figure 4D and 4E). In line with these observations, we also detected an increase in the ratio between oxidized versus reduced GSH (GSSH/GSH) in cKO BMDMs (Figure S7A-C). Given the central role of autophagy in mitochondrial turnover, we assessed mitochondrial morphology and respiration as a likely source of oxidative damage in uninfected cKO BMDMs (Figure S7D and S7E). However, no mitochondrial defect could be identified; this warrants further investigation into the underlying mechanism(s) of elevated oxidative stress in ATG16L1-deficient BMDMs.

Taken together, observations that cKO BMDMs are burdened with higher oxidative stress suggest that elevated ROS in these cells mandate upregulation of redox homeostasis factors in order to maintain viability (Tal et al., 2009). We thus asked whether accumulation of cytosolic ROS could be recapitulated by suppression of glutathione import in wild type macrophages. BMDMs were pre-treated with Erastin, a small molecule inhibitor of XCT, which diminishes the levels of reduced but not oxidized GSH in cells (Dixon et al., 2012). Time-course treatment demonstrated that cKO BMDMs are slightly more sensitive to Erastin after prolonged incubation, consistent with a basal elevation in cellular ROS (Figure S7F). Interestingly, 24-hour Erastin treatment of WT BMDMs phenocopied a steady-state ROS level in cKO cells, while no further increase in ROS was observed in cKO BMDMs treated with Erastin (Figure S7G). We hypothesized that induction of cellular ROS in WT BMDMs by pharmacological inhibition of XCT should phenocopy the accelerated *S.flexneri* clearance seen in cKO cells. To test this, WT BMDMs were pre-treated with Erastin for 18 hours prior infection and Erastin was maintained throughout the experiment (Figure 4F). Importantly, Erastin treatment did not increase WT BMDM cell death within the time-course (Figure S7H). However, Erastin-treated WT BMDMs showed enhanced elimination of *S.flexneri* following infection (Figure 4G and 4H), demonstrating that elevated oxidative stress in WT BMDMs accelerates killing of *S.flexneri*, consistent with enhanced microbicidal capacity of ATG16L1-deficient BMDMs.

## Discussion

Emerging insights from genetic mouse models have revealed that loss of *Atg16l1* in the immune and epithelial compartments lowers the threshold for an inflammatory response (Cadwell et al., 2010; Hubbard-Lucey et al., 2014; Lim et al., 2019; Matsuzawa-Ishimoto et al., 2017). Consistently, deletion of canonical autophagy genes in the innate and adaptive immune compartments have demonstrated enhanced pathogen clearance (Marchiando et al., 2013; Martin et al., 2018; Samie et al., 2018; Wang et al., 2020) as well as tumor control *in vivo* (Cunha et al., 2018; DeVorkin et al., 2019). These observations have prompted a re-evaluation of antimicrobial selective autophagy (xenophagy) to better understand how loss of core autophagy genes impacts cell-autonomous innate immunity against pathogenic intracellular bacteria.

In this study we show that macrophages deficient in ATG16L1 demonstrate an accelerated killing of *Shigella flexneri in vitro* and *in vivo*. To identify mechanisms behind this phenotype we employ isobaric multiplexing using the TMT technology, which emerged as being capable of near-comprehensive characterization of the global proteome (Lapek et al., 2017). When isobaric multiplexing methods are coupled with enrichment, it enables quantification of post-translational modifications on thousands of individual proteins (Rose et al., 2016). This method is ideally suited for interrogation of a complex response, such as infection of a host cell with an intracellular pathogen, where the diversity of downstream changes does not lend themselves to candidate approaches involving immunoblotting.

Our approach identifies multiple novel PTMs in components of inflammatory cytokine signaling, innate sensing and the core autophagy machinery that emerge as a consequence of *S.flexneri* infection. The comparison of early and late infection time-points shows a complex dynamic in the stability of PTMs as well as total protein abundance. The comparison of wild type versus ATG16L1-deficient BMDMs further reveals critical nodes in each of the above pathways that are under regulatory control by autophagy. The PTMs listed in Table 1, S2 and S3 represent a sizeable fraction of the relevant post-translational changes that occur in macrophages during infection and/or loss of autophagy. It is beyond the scope of a single study to interrogate these changes comprehensively; we encourage groups to utilize this study as a resource to explore PTMs in their pathway(s) of interest. We have provided interactive, web-accessible Spotfire Dashboards to enable user interrogation of the Global Proteome, Phospho-proteome, and the Ubiquitinome (KGG) datasets (https://info.perkinelmer.com/analytics-resource-center).

Our study reveals that basal accumulation of cellular ROS in cKO BMDMs enforces a compensatory increase in antioxidant responses exemplified by elevated protein abundances of key components of the glutathione synthesis machinery. This permits cellular viability under relatively elevated cytosolic ROS levels, which in turn suppresses *S.flexneri* expansion in BMDMs. However, overall macrophage fitness is likely compromised owing to a shift in the basal redox pathway set-point, and the accelerated clearance of *S.flexneri* observed in livers of *Atg16l1-cKO* mice may also contribute to inflammation-mediated loss of the hepatic myeloid cell niche *in vivo*. Pharmacological depletion of GSH phenocopies genetic loss of *Atg16l1*, accelerating *S.flexneri* clearance in wild type cells. These findings should prompt further investigation of autophagy in the intestinal epithelium, another key cellular niche for virulent *S.flexneri*. It is important to note that there are no viable murine models of enteric *S.flexneri* infection; development of model systems that permit intestinal infection while maintaining adequate inflammatory responses will be key in reconciling cell-based versus *in vivo* findings.

Our study provides the most comprehensive multiplexed proteomic analysis of the macrophage response to a cytosolic enteric pathogen to date. This novel resource will be of broad utility to the study of myeloid signal transduction, host-pathogen interaction and innate immunity.

## Materials and Methods

### Mice

All animal experiments were performed under protocols approved by the Genentech Institutional Animal Care and Use Committee (Protocol ID 17-2842). Generation of myeloid-specific deletion of *Atg16l1* was achieved by crossing *LysM*-Cre+ mice with *Atg16l1*^loxp/loxp^ mice and was described previously (Murthy et al., 2014). All mice were bred onto the C57BL/6N background. All *in vivo* experiments were performed using age-matched colony controls.

### Bacterial strains and culture

*Shigella flexneri* 5a strain M90T used in this study was obtained from ATCC (ATCC^®^ BAA-2402^™^). *Shigella flexneri* strain M90T Δ*mxiE* used in this study was obtained from a *S. flexneri* mutant collection (Sidik et al., 2014). Frozen bacterial stocks were streaked onto tryptic soy agar (TSA) plates and grown at 37 °C overnight. Plates were kept at 4 °C for up to 2 weeks.

### Bone marrow-derived cells isolation

Femurs and tibias were collected aseptically. After removing most of the muscle and fat, the epiphyses were cut and bones were placed into PCR tubes individually hung by the hinge into a 1.5 ml Eppendorf. The bone marrow was flushed by short centrifugation at 10,000 rpm for 30 seconds. Red blood cells were lysed with RBC lysis buffer (Genentech) by incubating for 5 minutes at RT. Cells were then pelleted and resuspended in BMDM media [high glucose Dulbecco’s Minimum Essential Media (DMEM) (Genentech) + 10% FBS (VRW, custom manufactured for Genentech) + GlutaMAX (Gibco, 30050-061) + Pen/Strep (Gibco, 15140-122) supplemented with 50 ng/ml recombinant murine macrophage-colony stimulating factor (rmM-CSF, Genentech)] and plated in 15-cm non-TC treated dishes for 5 days (Petri dish, VWR, 25384-326). Fresh BMDM media was added on day 3 without removal of original media. On day 5, macrophages were gently scraped from dishes, counted and re-plated on TC-treated plates of the desired format for downstream assays in fresh BMDM media. After overnight culture in BMDM media, assays were performed on day 6 BMDMs.

### BMDM infections in 24-well plates

BMDMs isolated from control *LysM-Cre+* or *LysM-Cre*+ Atg16L1^loxp/loxp^ mice were plated at 2.5 x 10^5^ cells/well in 24-well assay plates (Corning, 353047) in BMDM media. A duplicate plate was always plated for total PI-positive cell number enumeration after overnight incubation using IncuCyte ZOOM as described elsewhere. Bacterial cultures were prepared by picking a single bacterial colony from TSA plates and grown in 10 mL tryptic soy broth (TSB) in a shaking incubator overnight at 37 °C. After overnight incubation bacteria were subcultured in fresh 10 mL of TSB at 37 °C until OD600 0.5 - 0.8, pelleted by centrifugation, resuspended in 1:1000 poly-L-lysine (Sigma-Aldrich, P4707) in PBS and incubated for 10 minutes at RT. Cell suspension was then centrifuged and the pellet washed twice with PBS and once with the infection media [high glucose DMEM (Genentech) + 10% FBS (VRW, custom manufactured for Genentech) + GlutaMAX (Gibco, 30050-061)]. After the final wash the bacterial pellet was resuspended in the infection media and OD600 was remeasured. To prepare multiplicity of infection (MOI) of 5 in the infection media, total PI-positive object count per well was used for accurate MOI calculations for every independent infection experiment. A cell suspension containing lysine coated bacteria were added to the wells at MOI 5 in a total volume of 250 μl/well and allowed to adhere by incubating for 30 minutes at 37 °C in a CO2 incubator. After 30 minutes, bacterial suspension was aspirated and replaced with 500 μl/well of fresh infection media supplemented with gentamicin at 50 μg/mL (Sigma-Aldrich, G1397). This was defined as the time-point T = 0 minutes. Assay plates were subsequently incubated at 37 °C in a CO2 incubator and used at the indicated time-points for CFU enumeration.

### BMDM infections in 24-well plates with compounds

For experiments with Erastin (Sigma-Aldrich, E7781), day 5 BMDMs were plated at 2.5 x 10^5^ cells/well in 24-well assay plates (Corning, 353047) in BMDM media supplemented with Erastin at 4 μg/ml and incubated at 37 °C in a CO2 incubator for 18 hours before infection. A duplicate plate was also seeded and used for PI-positive object count per well enumeration to ensure accurate MOI as described elsewhere. The bacterial culture was prepared essentially as described elsewhere with the following modifications. After the final wash with infection media the bacterial pellet was resuspended in the infection media, OD600 was remeasured and bacterial suspension of MOI 10 was prepared. A cell suspension containing lysine coated bacteria was mixed 1:1 with infection media containing Erastin 8 μg/ml and added to the wells at MOI 5 in a total volume of 250 μl/well and allowed to adhere by incubating for 30 minutes at 37 °C in a CO2 incubator. After 30 minutes, bacterial suspension was aspirated and replaced with 500 μl/well of fresh infection media supplemented with gentamicin at 50 μg/mL (Sigma-Aldrich, G1397) and Erastin at 4 μg/ml as indicated.

### BMDM infections in 15-cm dishes for TMT proteomics

For large scale infections, 5-day differentiated BMDMs isolated from control *LysM-Cre*+ or *LysM-Cre*+ Atg16L1^loxp/loxp^ mice were plated at 10 x 10^6^ cells per 15-cm non-TC treated dish (Petri dish, VWR, 25384-326) in BMDM media. Bacterial suspension was prepared essentially as described elsewhere with the following modifications. A suspension of lysine coated bacteria in infection media were added to the dishes containing BMDMs at MOI 5 in a volume of 15 ml/dish and allowed to adhere by incubating for 30 minutes at 37 °C in a CO2 incubator. After 30 minutes, the medium was aspirated and replaced with 50 ml/dish of fresh infection media supplemented with gentamicin at 50 μg/mL (Sigma-Aldrich, G1397). This was defined as the time-point T = 0 minutes. Assay plates were subsequently placed at 37 °C in a CO2 incubator and samples collected after 30 - 45 minutes incubation (‘early’ infection time-point) or after 3 - 3.5 hours incubation (‘late’ infection time-point). At the indicated time-points a set of 10 dishes per genotype was used to prepare cell lysates for downstream proteomic analysis. To prepare cell lysates, infection media was first aspirated and cells washed once with PBS. Cells were then scrapped in the presence of Urea lysis buffer (20mM HEPES pH 8.0, 9M Urea, 1mM sodium orthovanadate, 2.5 mM sodium pyrophosphate, 1mM β-glycerolphosphate) and cell suspension stored at −80 °C until further processing(Kirkpatrick et al., 2013).

### *In vivo Shigella flexneri* infection

Mice were injected intravenously in the tail vein with *Shigella flexneri* (M90T) bacterial culture that was prepared by picking a single bacterial colony from TSA plates and grown in 10 mL tryptic soy broth (TSB) in a shaking incubator overnight at 37 °C. After overnight incubation bacteria were sub-cultured in fresh 10 mL of TSB at 37 °C until OD600 0.5 - 0.8, pelleted by centrifugation, washed with PBS once, resuspended in PBS and OD600 was recounted. Each animal was injected with 100 μl of bacterial suspension in PBS containing 2 x 10^6^ Colony Forming Units (CFUs) *S.flexneri (*M90T). Mice were euthanized after 6 or 24 hours post infection to harvest spleen and liver for CFUs enumeration and blood for cytokine profiling.

### Colony forming units (CFUs) assays

To determine CFUs in infected BMDMs, infection media was aspirated, cells were washed once with PBS and lysed by adding 250 μl/well of 0.1 % Igepal CA-630 (Sigma-Aldrich, I8896) in PBS, incubated for 5 minutes, resuspended and an aliquot of 200 μl was transferred to 96-well U-bottom plate (Costar, 3799) for making two-step serial dilutions in 0.1 % Igepal CA-630 in PBS. Subsequently, 5 μl of each serial dilution was plated on TSA plates in triplicates, allowed to evaporate at RT after which the plate was placed in a 37 °C incubator overnight. After overnight incubation, colonies from individual dilutions were counted and used for determining CFUs per well. To determine CFUs in the liver, mice were euthanized at the indicated time-points after infection and the livers were surgically removed and placed in PBS on ice. Livers were processed in 5 ml of 0.1 % Igepal CA-630 (Sigma-Aldrich, I8896) in PBS using the gentleMACS™ C Tubes (Miltenyi Biotec, 130-096-334) in combination with the gentleMACS™ Octo Dissociator (Miltenyi Biotec, 130-095-937) for the automated dissociation of tissues using standard tissue dissociation programs (program sequence: m_liver_01_02; m_liver_02_02, m_liver_01_02). Tissue suspensions were filtered through 100 μM filters (CellTreat, 229485) and remaining liver tissue was additionally homogenized using the rubber seal of the 5 ml syringe plunger. The resultant liver tissue suspension was used for generating serial dilutions and plated on TSA plates for CFUs enumeration as described elsewhere.

### IncuCyte assays

For IncuCyte assays, BMDMs were plated at 2 x 10^4^ cells/well in flat-bottom 96-well (Corning, 353072) or at 2.5 x 10^5^ cells/well in 24-well (Corning, 353047) assay plates. After overnight incubation at 37 °C in a CO2 incubator, cells were used for infection experiments or treatments with compounds or growth factors as indicated. BMDM viability over time was assessed by supplementing assay media [(high glucose DMEM (Genentech) + 10% FBS (VRW, custom manufactured for Genentech) + GlutaMAX (Gibco, 30050-061) + Pen/Strep (Gibco, 15140-122)] with propidium iodide (PI) dye for live-cell imaging at 1:1000 (Invitrogen, P3566), and then measuring PI-positive cells per mm^2^ using live cell imaging with IncuCyte ZOOM (IncuCyte systems, Essen Biosciences) in a time-course experiment. Percent cell death was calculated by dividing PI-positive cells per mm^2^ with total plated cells per mm^2^. Total plated cells were enumerated from a duplicate plate seeded at the same time as the assay plates. After overnight incubation, media in the duplicate plate was exchanged to assay media containing 0.06 % NP-40 supplemented with 1:1000 PI, and imaged at a single time-point using IncuCyte ZOOM after 10-minute incubation.

### GSH assays

BMDMs were established as described and 5×10^6 of BMDMs were pelleted by centrifugation, the pellet was lysed in mammalian lysis buffer (Abcam, ab179835), incubated 10’ at RT and centrifuged at top speed at 4°C 15min. Supernatant was transferred to a fresh tube and used for deproteinization following manufacturer’s instructions (Abcam, ab204708). The resultant supernatant was used for determining reduced GSH, total GSH and oxidized GSSG was calculated as per manufacturer’s instructions (Abcam, ab138881).

### Fluorescence microscopy

BMDMs grown on 96-well plates (Greiner Bio, 655090) were treated with 10 μM CellRox Green reagent for 30 minutes according to manufacturer’s protocol (Thermo Fisher Scientific, C10444), then fixed in 4 % paraformaldehyde (PFA) solution in PBS (ChemCruz, SC281692) for 15 minutes at RT. Nuclei were stained with NucBlue™ Live ReadyProbes™ Reagent (Thermo Fisher Scientific, R37605) for 10 minutes in PBS. 3D confocal images corresponding to 12 μm thick z-stacks of 4 stitched fields of views were collected on a Nikon A1R scanning confocal microscope using a Plan Apo NA 0.75 lens and x20 magnification. FITC and Hoechst 33342 signals were respectively imaged with the 488 nm and 405 nm laser lines. For each Z stack, images were combined into one focused image using Nikon Elements Extended Depth of focus (EDF) module that picks the focused regions from each frame and merges them together into a single focused image. The focused EDF images from different conditions were then analyzed with Bitplane Imaris software (version 9.2.0) using the cell segmentation module and intensity quantification. To specifically determine the cytoplasmic CellRox Green reagent intensity, the region corresponding to the Hoechst staining was excluded and FITC channel threshold was applied across all samples per given experiment. Mean cytosolic CellRox Green assay signal was then quantified per each individual cell and presented in the graph.

### Transmission Electron Microscopy

Samples were fixed in modified Karnovsky’s fixative (2% paraformaldehyde and 2.5% glutaraldehyde in 0.1M sodium cacodylate buffer, ph7.2) and then post-fixed in freshly prepared 1% aqueous potassium ferrocyanideosmium tetroxide (EM Sciences, Hatfield, PA), for 2h followed by overnight incubation in 0.5% Uranyl acetate at 4°C. The samples were then dehydrated through ascending series of ethanol (50%, 70%, 90%, 100%) followed by propylene oxide (each step was for 15 min) and embedded in Eponate 12 (Ted Pella, Redding, CA). Ultrathin sections (80 nm) were cut with an Ultracut microtome (Leica), stained with 0.2% lead citrate and examined in a JEOL JEM-1400 transmission electron microscope (TEM) at 80kV. Digital images were captured with a GATAN Ultrascan 1000 CCD camera.

## Tandem Mass Tag proteomics

### Protein Precipitation

Protein concentration in the lysates were quantified using the Pierce micro-BCA assay (ThermoFisher Scientific, Waltham, MA). All protein from the cell lysates was precipitated with a combination of methanol/chloroform/water(Wessel and Flügge, 1984). In brief, X volume of lysate was mixed with 4X volume of methanol followed by 2X volume of chloroform and 3X volume of water. The protein pellets were washed a total of three times with 5X volume of methanol. The protein pellets were air dried and resuspended in 8M urea, 100mM EPPS pH 7.0, 5mM DTT. Proteins were alkylated with 15 mM N-ethylmaleimide (Sigma).

### LysC/Trypsin Digestion

The protein in 8M urea was diluted to 4M with 100mM EPPS, pH 8.0. 15 mg of protein/sample was digested at 25 °C for 12 hours with lysyl endopeptidase (LysC, Wako Chemicals USA) at a 1:25; protein:protease ratio. Following LysC digestion the peptides in 4M urea were diluted to 1M urea with 100mM EPPS, pH 8.0. The LysC peptides were digested with trypsin at 37 °C for 8 hours (Promega) at a 1:50; protein:protease ratio.

### Ubiquitin Remnant Peptide Enrichment (KGG peptides)

Prior to KGG peptide enrichment, the tryptic peptides were acidified to 2% formic acid and desalted with 1 g tC18 Sep-Pak cartridges (Waters). The desalted peptides were dried by vacuum. KGG peptide enrichment was performed with the PTMScan ubiquitin remnant motif kit (Cell Signaling Technologies, Kit#5562) as per the manufacturers protocol. KGG peptides eluted from the antibodies were dried by vacuum. The flow through peptides from the KGG enrichment were saved for phosphopeptide and total protein analysis.

### TMT labelling of KGG Peptides

Peptides were resuspended in 200mM EPPS, pH 8.0. 10 μL of TMT reagent at 20 μg/uL (ThermoFisher) was added to each sample. Peptides were incubated with TMT reagent for 3 hours at 25 °C. TMT-labeled peptides were quenched with hydroxylamine (0.5% final) and acidified with trifluoroacetic acid (2% final). The samples were combined, desalted with 50 mg tC18 Sep-Paks, and dried by vacuum.

### Ubiquitin Remnant Peptide Fractionation

TMT-labeled KGG peptides were fractionated using the high pH reversed-phase peptide fractionation kit (ThermoFisher). The dried KGG peptides were resuspended in 0.1% trifluoroacetic acid and fractionated according to the manufacturers protocol into 6 fractions (17.5%, 20%, 22.5%, 25%, 30%, and 70% acetonitrile + 0.1% triethylamine). The KGG peptide fractions were dried by vacuum, desalted with StageTips packed with Empore C18 material (3M, Maplewood, MN.), and dried again by vacuum. KGG peptides were reconstituted in 5% formic acid + 5% acetonitrile for LC-MS3 analysis.

### TMT labelling of KGG Flow Through Peptides

The flow through peptides from the KGG enrichment were labeled with TMT prior to phosphopeptide enrichment. The flow through peptides were resuspended in 1X IAP buffer from the ubiquitin remnant kit (from prior step). The pH of the resuspended peptides was adjusted by adding 1M EPPS, pH 8.0 in a 3:1 ratio (peptide volume:1M EPPS volume; 250mM EPPS final). 2.1 mg of peptide from each sample was labeled with 2.4 mg of TMT reagent resuspended in 60 μL, 100% acetonitrile. The peptides were incubated with TMT reagent for 3 hours at 25 °C. TMT-labeled peptides were quenched with hydroxylamine (0.5% final) and acidified with trifluoroacetic acid (2% final). The samples were combined, desalted with 1 g tC18 Sep-Paks, dried by vacuum.

### Phosphoserine, -threonine, -tyrosine Enrichment and Fractionation

Phosphotyrosine (pY) peptides were enriched using the Cell Signaling Technologies pY-1000 antibody kit as per the manufacturers protocol (Cell Signaling Technologies, Kit#8803). The flow through from the pY enrichment was desalted on a 1g tC18 Sep-Pak cartridge (Waters Corporation, Milford, MA) and dried by centrifugal evaporation and saved for phosphoserine and phosphothreonine (pST) analysis. pST phosphopeptides were enriched using the Pierce Fe-NTA phospho-enrichment kit (ThermoFisher). In brief, peptides were bound and washed as per manufacturers protocol. Phosphopeptides were eluted from the Fe-NTA resin with 50mM HK2PO4 pH 10.5. Labelled phosphopeptides were subjected to orthogonal basic-pH reverse phase fractionation on a 3×100 mm column packed with 1.9 μm Poroshell C18 material (Agilent, Santa Clara, CA), utilizing a 45 min linear gradient from 8% buffer A (5% acetonitrile in 10 mM ammonium bicarbonate, pH 8) to 30% buffer B (acetonitrile in 10mM ammonium bicarbonate, pH 8) at a flow rate of 0.4 ml/min. Ninety-six fractions were consolidated into 18 samples, acidified with formic acid and vacuum dried. The samples were resuspended in 0.1% trifluoroacetic acid, desalted on StageTips and vacuum dried. Peptides were reconstituted in 5% formic acid + 5% acetonitrile for LC-MS3 analysis. The flow-through peptides from the pST enrichment were saved for total protein analysis.

### Peptide Fractionation for Total Protein Analysis

The flow-through from the pST enrichment was dried by centrifugal evaporation. The dried peptides were resuspended in 0.1% TFA. Approximately 250 μg of peptide mix was subjected to orthogonal basic-pH reverse phase fractionation on a 3×100 mm column packed with 1.9 μm Poroshell C18 material (Agilent, Santa Clara, CA), utilizing a 45 min linear gradient from 8% buffer A (5% acetonitrile in 10 mM ammonium bicarbonate, pH 8) to 35% buffer B (acetonitrile in 10mM ammonium bicarbonate, pH 8) at a flow rate of 0.4 ml/min. Ninety-six fractions were consolidated into 12 samples, acidified with formic acid and vacuum dried. The samples were resuspended in 5% formic acid, desalted on StageTips and vacuum dried. Peptides were reconstituted in 5% formic acid + 5% acetonitrile for LC-MS3 analysis.

### Mass spectrometry analysis

All mass spectra were acquired on an Orbitrap Fusion Lumos coupled to an EASY nanoLC-1000 (or nanoLC-1200) (ThermoFisher) liquid chromatography system. Approximately 2 μg of peptides were loaded on a 75 μm capillary column packed in-house with Sepax GP-C18 resin (1.8 μm, 150 Å, Sepax Technologies) to a final length of 35 cm. Peptides for total protein analysis were separated using a 180-minute linear gradient from 8% to 23% acetonitrile in 0.1% formic acid. The mass spectrometer was operated in a data dependent mode. The scan sequence began with FTMS1 spectra (resolution = 120,000; mass range of 350-1400 *m/z*; max injection time of 50 ms; AGC target of 1e6; dynamic exclusion for 60 seconds with a +/- 10 ppm window). The ten most intense precursor ions were selected for ITMS2 analysis via collisional-induced dissociation (CID) in the ion trap (normalized collision energy (NCE) = 35; max injection time = 100ms; isolation window of 0.7 Da; AGC target of 2e4). Following ITMS2 acquisition, a synchronous-precursor-selection (SPS) MS3 spectrum was acquired by selecting and isolating up to 10 MS2 product ions for additional fragmentation via high energy collisional-induced dissociation (HCD) with analysis in the Orbitrap (NCE = 55; resolution = 50,000; max injection time = 110 ms; AGC target of 1.5e5; isolation window at 1.2 Da for +2 *m/z*, 1.0 Da for +3 *m/z* or 0.8 Da for +4 to +6 *m/z*).

pY peptides were separated using a 180-minute linear gradient from 7% to 26% acetonitrile in 0.1% formic acid. The mass spectrometer was operated in a data dependent mode. The scan sequence began with FTMS1 spectra (resolution = 120,000; mass range of 350-1400 *m/z*; max injection time of 50 ms; AGC target of 1e6; dynamic exclusion for 75 seconds with a +/- 10 ppm window). The ten most intense precursor ions were selected for FTMS2 analysis via collisional-induced dissociation (CID) in the ion trap (normalized collision energy (NCE) = 35; max injection time = 150ms; isolation window of 0.7 Da; AGC target of 3e4; *m/z* = 2-6; Orbitrap resolution = 15k). Following FTMS2 acquisition, a synchronous-precursor-selection (SPS) MS3 method was enabled to select five MS2 product ions for high energy collisional-induced dissociation (HCD) with analysis in the Orbitrap (NCE = 55; resolution = 50,000; max injection time = 300 ms; AGC target of 1e5; isolation window at 1.2 Da.

pST peptides were separated using a 120-minute linear gradient from 6% to 26% acetonitrile in 0.1% formic acid. The mass spectrometer was operated in a data dependent mode. The scan sequence began with FTMS1 spectra (resolution = 120,000; mass range of 350-1400 *m/z*; max injection time of 50 ms; AGC target of 1e6; dynamic exclusion for 60 seconds with a +/- 10 ppm window). The ten most intense precursor ions were selected for ITMS2 analysis via collisional-induced dissociation (CID) in the ion trap (normalized collision energy (NCE) = 35; max injection time = 200ms; isolation window of 0.7 Da; AGC target of 2e4). Following MS2 acquisition, a synchronous-precursor-selection (SPS) MS3 method was enabled to select five MS2 product ions for high energy collisional-induced dissociation (HCD) with analysis in the Orbitrap (NCE = 55; resolution = 50,000; max injection time = 300 ms; AGC target of 1e5; isolation window at 1.2 Da for +2 *m/z*, 1.0 Da for +3 *m/z* or 0.8 Da for +4 to +6 *m/z*).

KGG peptides were separated using a 180-minute linear gradient from 7% to 24% acetonitrile in 0.1% formic acid. The mass spectrometer was operated in a data dependent mode. The scan sequence began with FTMS1 spectra (resolution = 120,000; mass range of 350-1400 *m/z*; max injection time of 50 ms; AGC target of 1e6; dynamic exclusion for 75 seconds with a +/- 10 ppm window). The ten most intense precursor ions were selected for FTMS2 analysis via collisional-induced dissociation (CID) in the ion trap (normalized collision energy (NCE) = 35; max injection time = 100ms; isolation window of 0.7 Da; AGC target of 5e4; *m/z* 3-6, Orbitrap resolution set to 15k). Following MS2 acquisition, a synchronous-precursor-selection (SPS) MS3 method was enabled to select 10 MS2 product ions for high energy collisional-induced dissociation (HCD) with analysis in the Orbitrap (NCE = 55; resolution = 50,000; max injection time = 500 ms; AGC target of 1e5; isolation window at 1.0 Da for +3 *m/z* or 0.8 Da for +4 to +6 *m/z*).

MS/MS spectra for the global proteome, serine/threonine phosphorylated, tyrosine phosphorylated, and ubiquitylated data sets were searched using the Mascot search algorithm (Matrix Sciences) against a concatenated target-decoy database comprised of the UniProt mouse and *Shigella flexneri* protein sequences (version 2017_08), known contaminants and the reversed versions of each sequence. For all datasets a 50 ppm precursor ion mass tolerance was selected with tryptic specificity up to two missed cleavages. For the global proteome and serine/ threonine phosphorylated datasets a 0.8 Da fragment ion tolerance was selected. While for the tyrosine phosphorylated and KGG (ubiquitin) datasets a 0.02 Da fragment ion tolerance was selected. The global proteome and phosphorylated datasets used a fixed modification of N-ethylmaleimide on cysteine residues (+125.0477) as well as TMT 11-plex on Lysine and the peptide N-term (+229.1629). The ubiquitylated data set used a fixed modification of N-ethylmaleimide on cysteine residues (+125.0477) as well as TMT 11-plex on the peptide N-term (+229.1629). For variable modifications the global proteome dataset used methionine oxidation (+15.9949) as well as TMT 11-plex on tyrosine (+229.1629). The phosphorylated dataset used the same variable modifications as the global proteome dataset plus phosphorylation on serine, threonine, and tyrosine (+79.9663). Finally, the ubiquitylated dataset used methionine oxidation (+15.9949), TMT 11 plex on tyrosine and lysine (+229.1629), as well as TMT 11 Plex + ubiquitylation on lysine (343.2059). PSMs were filtered to a 1% peptide FDR at the run level using linear discriminant analysis (LDA) (Kirkpatrick et al., 2013). PSM data within each plex and dataset (global proteome, phosphorylation, and ubiquitylation) was aggregated and these results were subsequently filtered to 2% protein FDR. For PSMs passing the peptide and protein FDR filters within the phosphorylated and ubiquitylated datasets, phosphorylation and ubiquitylation site localization was assessed using a modified version of the AScore algorithm(Beausoleil et al., 2006) and reassigned accordingly. Finally, reporter ion intensity values were determined for each dataset and plex using the Mojave algorithm(Zhuang et al., 2013) with an isolation width of 0.7.

### Quantification and statistical testing of global proteomics and phosphoproteomic data

Quantification and statistical testing of global proteomics data were performed by MSstatsTMT v1.2.7, an open-source R/Bioconductor package(Huang et al., 2020; Tsai et al., 2020). MSstatsTMT was used to create quantification reports and statistical testing reports using the Peptide Spectrum Matches (PSM) as described above. First, PSMs were filtered out if they were (1) from decoy proteins; (2) from peptides with length less than 7; (3) with isolation specificity less than 70%; (4) with reporter ion intensity less than 2^8 noise estimate; (5) from peptides shared by more than one protein; (6) with summed reporter ion intensity (across all eleven channels) lower than 30,000; (7) with missing values in more than nine channels. In the case of redundant PSMs (i.e., multiple PSMs in one MS run corresponding to the same peptide ion), only the single PSM with the least missing values or highest isolation specificity or highest maximal reporter ion intensity was retained for subsequent analysis. Multiple fractions from the same TMT mixture were combined in MSstatsTMT. In particular, if the same peptide ion was identified in multiple fractions, only the single fraction with the highest mean or maximal reporter ion intensity was kept. Next, MSstatsTMT generated a normalized quantification report across all the samples at the protein level from the processed PSM report. Global median normalization, which equalized the median of the reporter ion intensities across all the channels and TMT mixtures, was carried out to reduce the systematic bias between channels. The normalized reporter ion intensities of all the peptide ions mapped to a protein were summarized into a single protein level intensity in each channel and TMT mixture. For each protein, additional local normalization on the summaries was performed to reduce the systematic bias between different TMT mixtures. For the local normalization, we created an artifact reference channel by averaging over all the channels except 131C for each protein and TMT mixture. The channel 131C was removed in order to make each mixture have the same number of samples from each condition. The normalized quantification report at the protein level is available in Supplementary Table 6. As a final step, the differential abundance analysis between conditions was performed in MSstatsTMT based on a linear mixed-effects model per protein. The inference procedure was adjusted by applying an empirical Bayes shrinkage. The table with the statistical testing results for all the proteins is available as in Supplementary Table 7. Quantification and statistical testing for phospho- and KGG (Ub) site data were performed by the same procedure as for global proteomics data with some modifications. First, PSMs from non-modified peptides were filtered out from the PSM report and the remaining preprocessing analyses were the same as above. Second, custom PTM site identifiers were created for each PSM by identifying the modified residue index in the reference proteome that was used to search the MS/MS spectra. Finally, all steps for quantification and differential abundance analysis were performed at the PTM site level, rather than the protein level (Supplementary Tables 8 and 9). The relative abundance of TMT reporter ion abundances in bar graphs throughout the paper stems from MSstats modeling and sums up to 1.0 for each Plex. Thus, the sum of all signal shown sums to 1.0 or 2.0 depending on whether the feature was quantified in one or both plexes. For the consolidated heatmaps showing proteome level changes immediately adjacent to any identified PTMs, the ComplexHeatmap R package was used.

### Gene set enrichment analysis

Gene set enrichment analysis was performed using MsigDB(Liberzon et al., 2015; Subramanian et al., 2005). Global proteome data were filtered to include features with an absolute value log2fc values of greater than 1 as well as p values of less than 0.05. Subsequently the data were filtered to require that every protein must be found in both multiplexed experiments. UniProt identifiers were transformed to gene symbols and fed into GSEA for an enrichment analysis against MsigDB’s hallmark gene sets. Gene set enrichment results were filtered to 5% FDR.

### Overview Heatmaps/Clustering

For the overview heatmaps showing PTM and global proteome datasets side by side, clustering was performed as follows. First, protein quantification results from MSstatsTMT for the PTM and global proteome datasets were merged with the phospho-proteome and KGG datasets, respectively. For each of the two combined datasets, the pheatmap R package was used to cluster the protein model results into 16 row wise clusters using the clustering method ‘ward.D’. The columns of the dataset were kept static and not clustered.

### Statistical analysis

Pairwise statistical analyses were performed using an unpaired t-test using two-stage step-up method of Benjamini, Krieger and Yekutieli and false discovery rate of 1% to determine if the values in two sets of data differ. Multiple-comparison corrections were made using the Sidak method with family-wise significance and confidence level of 0.05. Analysis of *in vivo* infection data was done using unpaired two-tailed t-test after outliers were removed using ROUT method (Q = 1 %). Analysis of kinetic (time) with Erastin was performed using two-way ANOVA followed by multiple comparison testing. Line graphs and associated data points represent means of data; error bars represent standard deviation from mean. GraphPad Prism 8 software was used for data analysis and representation. P-values: *<0.05, **<0.01, ***<0.001, ****<0.0001. For proteomics data, differential abundance analysis between conditions and p-values were determined based on a linear mixed-effects model per protein (global proteome data) or per PTM site (Phosphorylation, Ubiquitin-KGG data) using MSstatsTMT software package.

## Data availability

Mass spectrometry raw files have been uploaded to the UCSD MassIVE repository and are available: (https://massive.ucsd.edu/ProteoSAFe/dataset.jsp?accession=MSV000085565; Password= shigella).

## Software Availability

Raw files were converted to mzXML using ReadW (v 4.3.1) available through https://sourceforge.net/projects/sashimi/files/ReAdW%20%28Xcalibur%20converter%29/. Spectra were searched using Mascot (v 2.4.1) licensed from Matrix Sciences. Search results were filtered using the LDA function in the MASS Package in R as described in Huttlin et al. Cell 143, 1147-1189 (2010). Mojave is an in-house tool developed to report TMT reporter ion intensity values and is available upon request. MSstatsTMT (v 1.2.7) is a freely available open-source R/Bioconductor package to detect differentially abundant proteins in TMT experiments. It can be installed through https://www.bioconductor.org/packages/release/bioc/html/MSstatsTMT.html. Gene set enrichment was performed using the GSEA/MSigDB web portal https://www.gsea-msigdb.org/gsea/msigdb/annotate.jsp. Heatmaps were generated using the pheatmap (v1.0.12) (https://cran.r-project.org/web/packages/pheatmap/index.html) or ComplexHeatmap (v 2.4.2) (https://bioconductor.org/packages/release/bioc/html/ComplexHeatmap.html) R packages.

## Acknowledgements

We thank Avinashnarayan Venkatanarayan and the laboratory of Eric Brown at Genentech for technical assistance.

## Author contributions

T.M., I.D., D.S.K. and A.M. designed the conceptual framework of the study and experiments. T.M. designed and performed large-scale proteomic experiments with assistance from J.L. and guidance from I.D., D.S.K. and A.M. TMT data acquisition and initial data analysis performed by R.C.K., B.K.E., T.H., M.C., T-H. T. and O.V. TMT data analysis and representation performed by T.H., E.V. and D.S.K. with input from T.M. and A.M. Electron microscopy performed by A.K.K. and M.R. CellRox microscopy experiments performed by T.M., P.C. and C.C. *In vitro* BMDM infection assays performed by T.M. *In vivo* infection experiments performed by T.M. and A.M. J.R. provided bacterial strains and guided the infection studies. The manuscript was written by T.M. D.S.K. and A.M with contributions and comments from all authors.

## Supplemental Figures and Tables

**Supplemental Figure 1.**
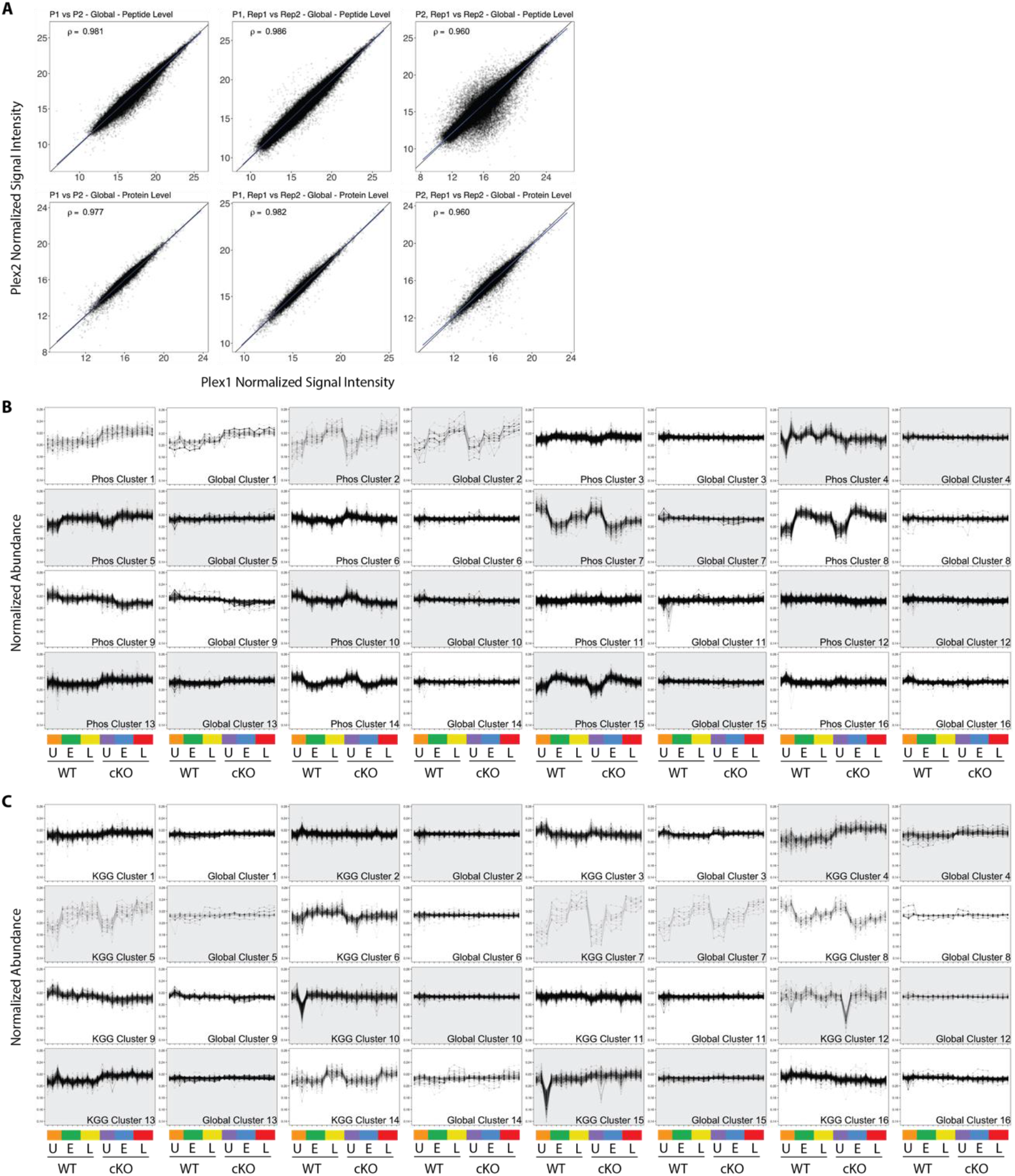
Quality control and PTM-Global comparative analysis of proteomics data. (A) Scatterplots showing normalized signal intensity from the Global Proteome analysis. Peptide (upper row) and protein (lower row) level data stemming from MSstats modeling are displayed. Plots in the first column compare Ple× 1 versus Plex 2, where data from intra-plex duplicates was aggregated during modeling. Plots in the middle and right columns compare intra-plex duplicate samples within Plexl (middle) or Plex2 (right). Pearson correlations are shown for each contrast. (B and C) Line plots showing all 16 K-means clusters corresponding to the Phospho-Global (B) and KGG (Ub)-Global (C) heatmaps displayed in Figure 2. The background shading for each pair of line plots is toggled to highlight pairing between Phos/KGG and Global protein clusters.

**Supplemental Figure 2.**
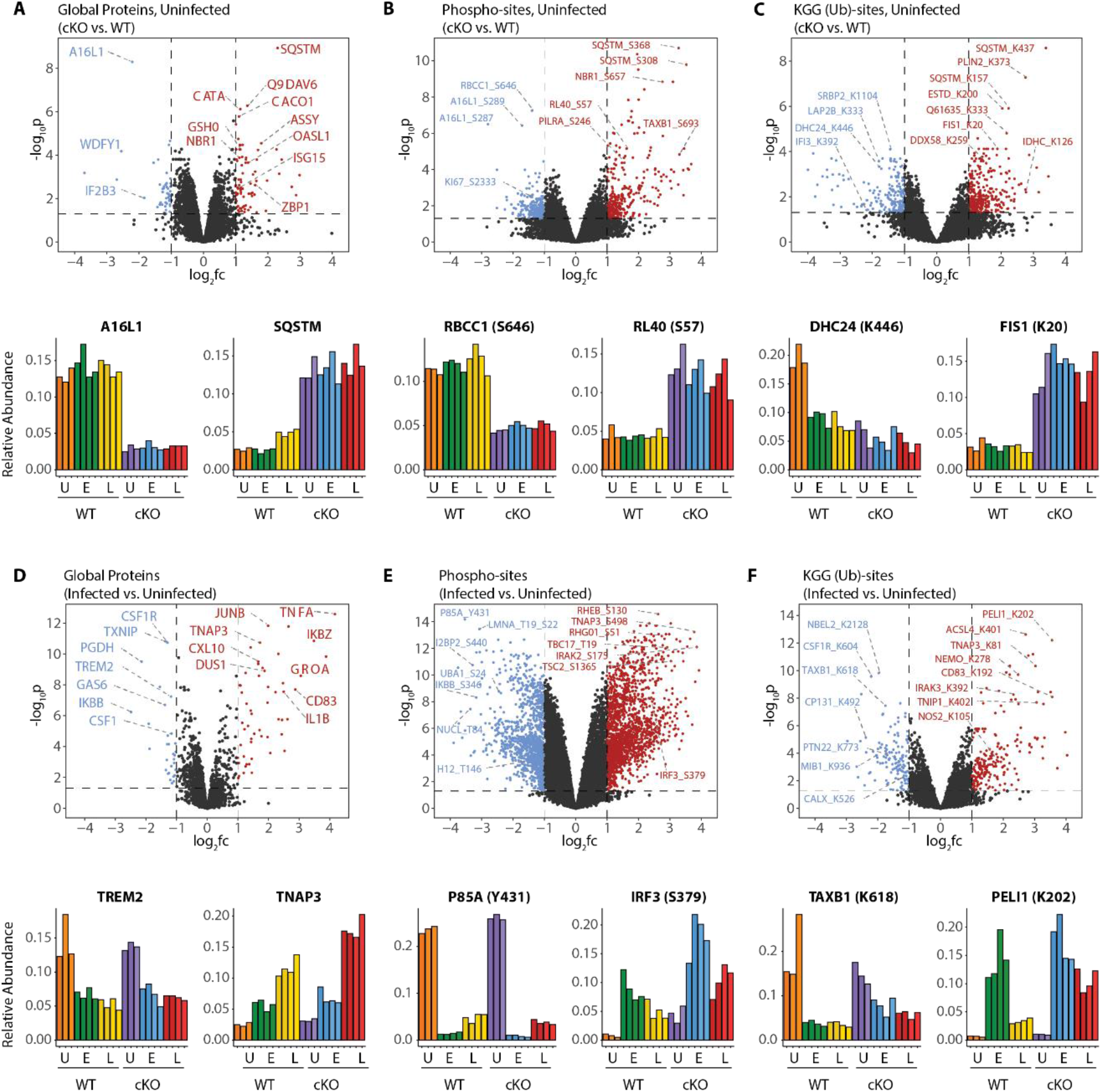
A global overview of changes identified between the genotypes and upon infection. (A-C) Volcano plots showing differential expression of Global Proteins (A), Phospho-sites (B) and KGG (Ub)-sites (C) between uninfected cKO vs. WT BMDMs. Volcano plots display log2 fold changes and -log_10_ transformed p-values for the host proteome. Bar graphs at the bottom of each panel represent top hits with positive and negative log2 fold changes. Uninfected (U) samples are shown with orange (WT) and purple (cKO), early infection (E) in green (WT) and blue (cKO) and late infection in yellow (WT) and red (cKO), respectively. Protein names are shown as UniProt identifiers with modification sites indicated by the modified amino acid (S/T/Y/K) and residue number (e.g. RL4Ũ_S57). Features enriched in cKO and WT BMDMs are highlighted in red and blue, respectively. (D-F) Volcano plots displaying differentially expressed Global Proteins (D), Phospho-sites (E) and KGG (Ub)-sites (F) between infected and uninfected BMDMs. Infected refers to the aggregate condition in which early (E) and late (L) infected samples for WT and cKO are each weighted as 0.25 relative to 0.5 each for the WT and cKO uninfected samples. Features enriched in infected and uninfected BMDMs are highlighted in red and blue, respectively. As above, bar graphs below each panel show example hits. The relative abundance of TMT reporter ions sums up to 2.0 for features quantified in both Plexl and Plex2.

**Supplemental Figure 3.**
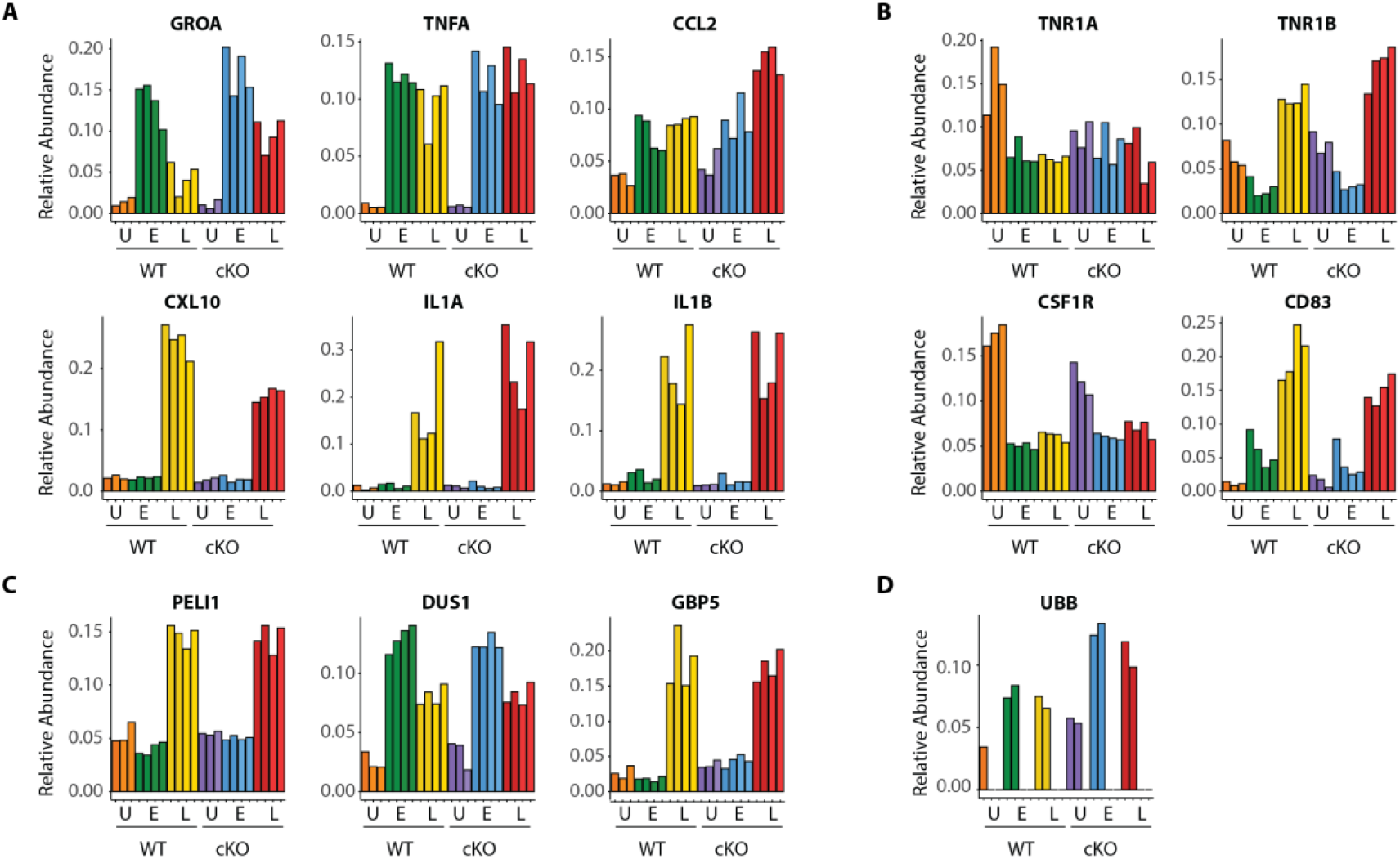
Dynamic macrophage response to infection. (A-D) Bar graphs showing quantitative changes for representative pro-inflammatory cytokines and chemokines (A), cell surface receptors (B), components of innate immune signaling (C) and linear ubiquitin chains as represented by UBB (D).

**Supplemental Figure 4.**
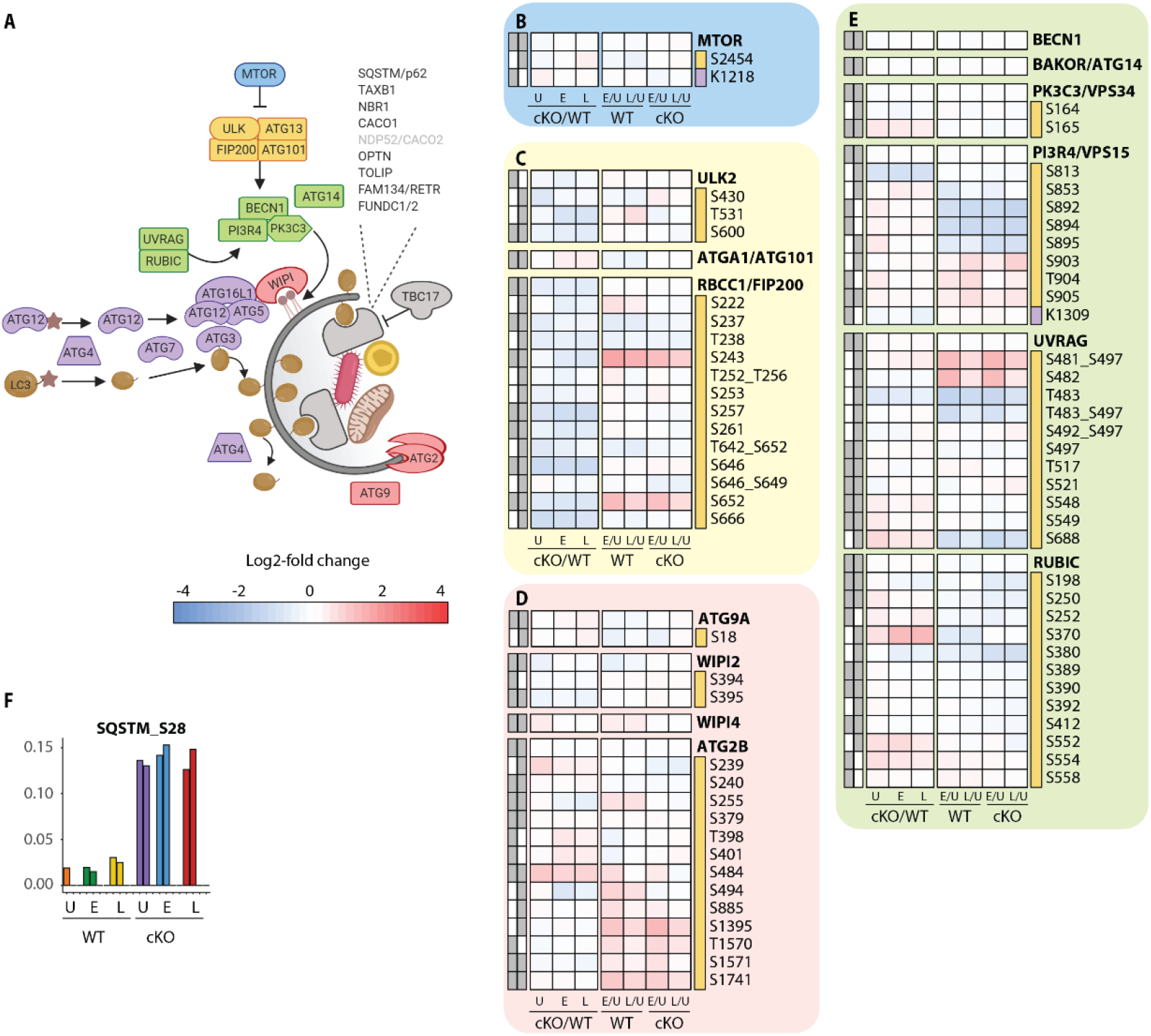
Extended analysis of proteomic changes in the autophagy pathway. (A) Schematic representation of macro-autophagy & selective autophagy machinery as shown in Figure 3C. (B-E) Heatmap representations of mTOR (B), ULK complex (C), membrane recruitment and closure (D), and PIK3C3Λ/ps34 complexes (E) are shown. The background shading for each panel corresponds to the functional color coding of proteins in the pathway schematic shown in (A). (F) Bar graph showing phosphorylation on autophagy receptor p62 (SQSTM_S28).

**Supplemental Figure 5.**
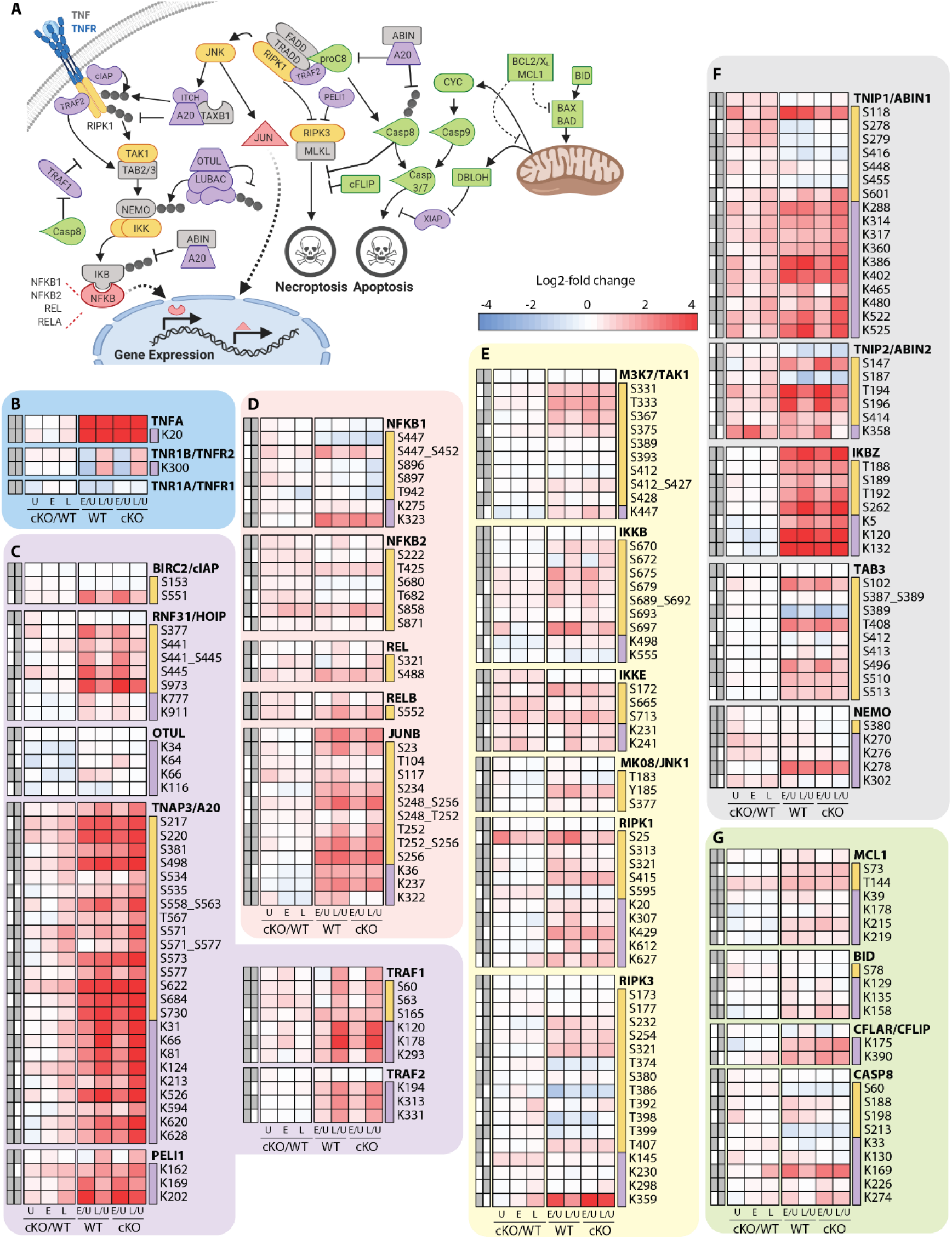
Characterization of proteomic changes in inflammatory signaling nodes. (A) Schematic representation of key components in the inflammatory signaling and programmed cell death pathways analyzed in this study (B-G) Heatmap representations of TNF and its receptors (B), E3 ubiquitin ligase and deubiquitinase enzymes (C) and transcription factors (C), kinases (E), signaling adaptors (F), and apoptosis regulatory proteins (G). The background shading for each panel corresponds to the functional color coding of proteins in the pathway schematic shown in (A).

**Supplemental Figure 6.**
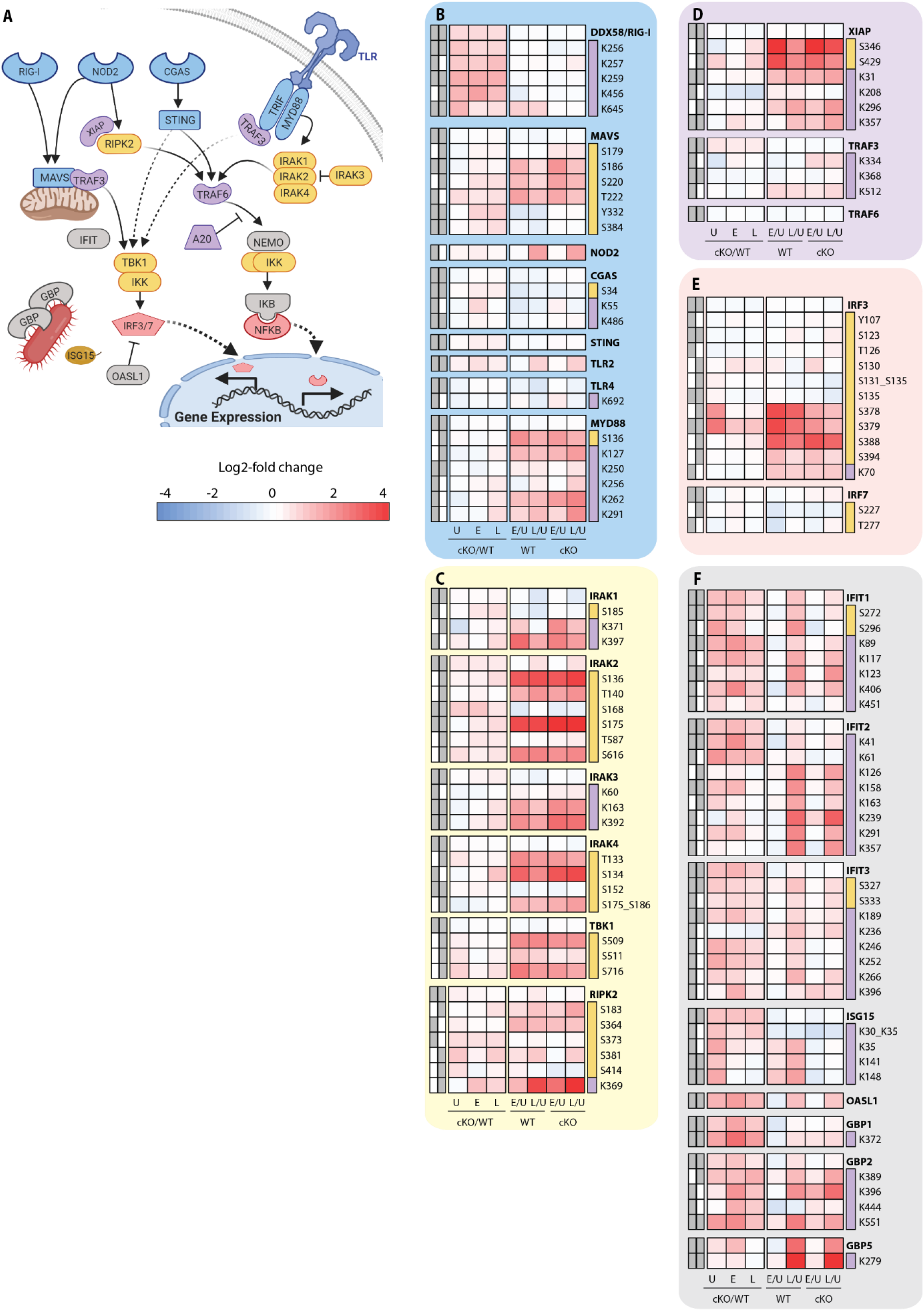
Analysis of proteomic changes in innate sensing and the interferon response. (A) Schematic representation of innate sensors and inflammatory signaling components analyzed in this study. (B-F) Heatmap representations of microbe-associated molecular pattern receptors and adaptors (B), kinases (C), E3 ubiquitin ligases (D), transcription factors (E) and interferon response genes (F) are shown. The background shading for each panel corresponds to the functional color coding of proteins in the pathway schematic shown in (A).

**Supplemental Figure 7.**
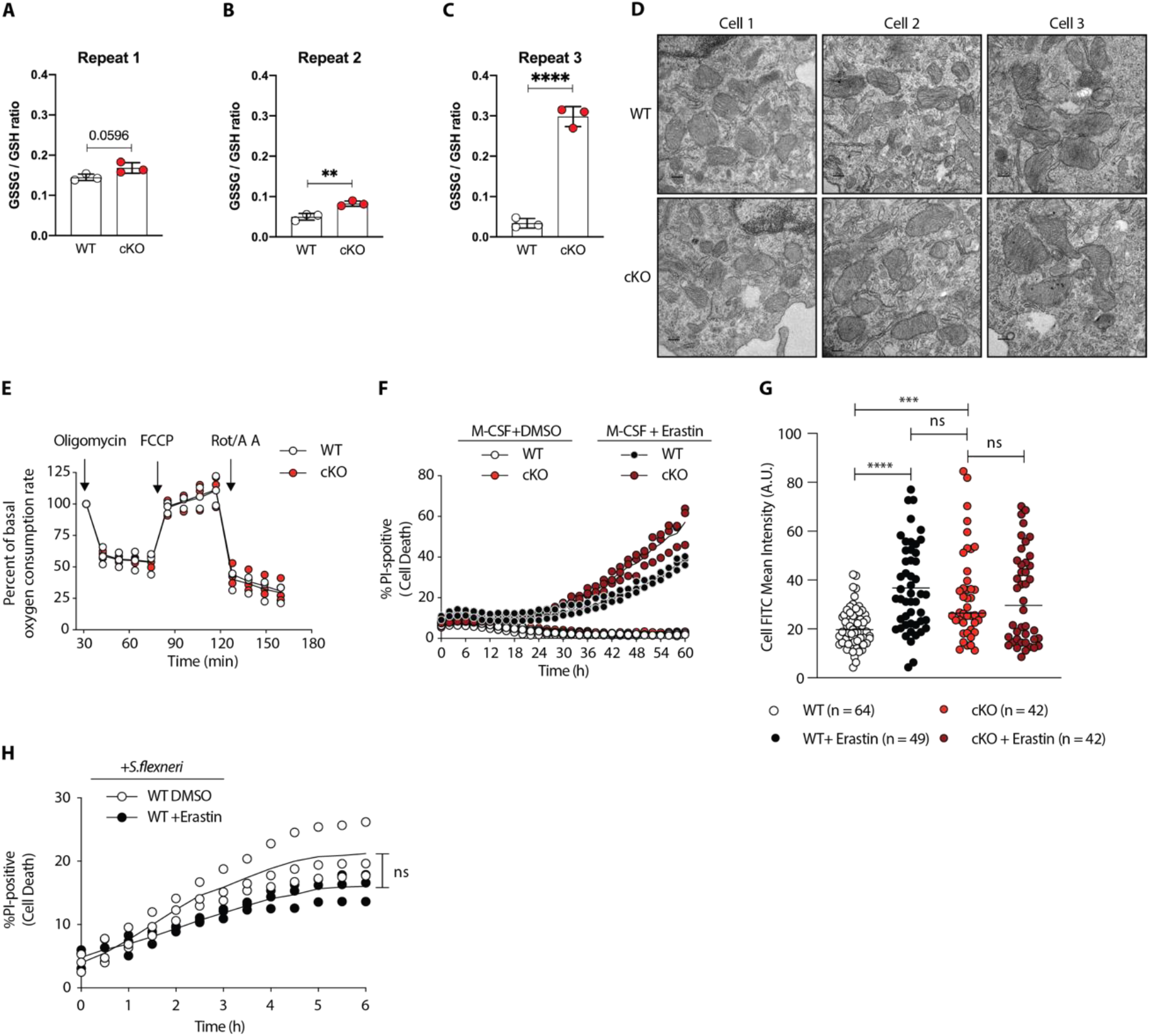
Elevated oxidative stress in ATG16L1-deficient macrophages. (A-C) Graphs show three independent experiments quantifying the GSSG/GSH ratio in BMDMs from three different preparations. Unpaired t test “ P = 0.0057, **** P < 0.0001. (D) Representative electron microscopy (EM) micrographs of WT and cKO BMDMs at magnification 8000x (scale bar = 0.2 μm). (E) Seahorse assay showing oxygen consumption rate of WT and cKO BMDMs. Graph shows the percent of basal respiration following treatment with 1.5 μM oligomycin, 1 μM FCCP and 0.5 μM Rotenone/Antimycin A from three independent experiments using three different BMDM preparations (n = 3). (F) Percentage of Pl-positive WT and cKO BMDMs during time-course incubation with DMSO or 4 μg/ml Erastin. A representative graph shows individual values from three wells from one experiment (n = 3). (G) Quantification of CellRox green probe mean intensity in WT and cKO BMDMs in the absence or presence of Erastin 4 μg/ml for 24h. Graph shows single cell data from one experiment (n = 3). Ordinary one-way ANOVATukey’s multiple comparison test **** P < 0.0001 and *** P = 0.0003. Apart of the data is also used in Figure 4E. (H) Percentage of Pl-positive WT BMDMs during time-course infection with *S.flexneri* M9OT in the presence of DMSO or Erastin 4 μg/ml. Graph represents individual values from three independent experiments using three different BMDM preparations, ns, non-significant.

**Supplemental Table 1. Composition PTM-Site and Global Protein clusters displayed in Figure 1F, 1G and S1b, S1C.**

**Supplemental Table 2. Curated list of PTMs described in Figure 3 and S3 with associated references.**

**Supplemental Table 3. Curated list of PTMs described in Figure 4 and S4 with associated references.**

**Supplemental Table 4. Gene Set Enrichment Analysis (GSEA) performed to identify cellular processes overrepresented in ATG16L1 deficient BMDMs in Figure 5A.**

**Supplemental Table 5. Protein references and gene names associated with mitochondrial and peroxisomal categories in Figure S5A-C.**

**Supplemental Table 6. MSstatsTMT normalized quantification report for Global Proteins data.**

**Supplemental Table 7. MSstatsTMT statistical testing results for Global Proteins data.**

**Supplemental Table 8. MSstatsTMT normalized quantification report for Phosphorylation Site data.**

**Supplemental Table 9. MSstatsTMT normalized quantification report for KGG (Ub)-sites data.**

